# Phosphatidylinositol-3,5-bisphosphate mediated vacuolar morphology modulation is integral to ethanol stress response

**DOI:** 10.1101/2024.12.19.629339

**Authors:** Pritha Mandal, Babai Hazra, Trisha Ghosh, Nabanita Patra, Srimonti Sarkar

## Abstract

Vacuoles enlarge in response to ethanol. We used quantitative imaging to show that even brief exposure to ethanol causes a rapid transition from multilobed vacuoles to a single compartment. Vacuole unlobing is essential as mutants restricted for vacuole fusion exhibit greater ethanol sensitivity. This response involves the inhibition of vacuole fission via downregulation of the activity of Fab1, a lipid kinase generating the signaling lipid phosphatidylinositol-3,5-bisphosphate. Ethanol exposure results in the redistribution of a phosphatidylinositol-3,5-bisphosphate sensor from the perivauolar dots to the cytoplasm and the dissociation of the effector of this lipid, Atg18, from the vacuolar membrane to the cytoplasm. Impaired Atg18 membrane recruitment also entails a loss of interaction between a scaffold of the Fab1 complex, Vac14, and Atg18. A hyperactive Fab1 mutant, with elevated pools of phosphatidylinositol-3,5-bisphosphate, shows delayed vacuole enlargement; the partial relocation of the phosphatidylinositol-3,5-bisphosphate probe to the perivacuolar dots and of Atg18 to the vacuole membrane. Since Atg18 is responsible for vacuole fragmentation, our results show that, under ethanol stress, cells actively regulate vacuole morphology by modulating phosphatidylinositol-3,5-bisphosphate levels.

## INTRODUCTION

Bioethanol is a key renewable energy resource. A major avenue of bioethanol production is via the fermentation of carbohydrates using yeast *Saccharomyces cerevisiae*. Ethanol yield from fermentation is limited by the toxic effect of this fermentation product on yeast. Ethanol’s toxicity is manifested at multiple levels and includes i) disruption of membrane integrity (Goldstein, 1986), ii) increased cytoplasmic acidification due to the entry of ions, especially protons, through breaches in the plasma membrane (Auesukaree, 2017), and iii) structural destabilization of cellular proteins (Feng *et al*., 2021). As fermentation proceeds, the concentration of ethanol increases in the growth medium, which impacts both the growth and viability of yeast, ultimately resulting in the cessation of ethanol production (Carmona-Gutierrez *et al*., 2012). To increase ethanol yield, an understanding of the cellular pathways used by yeast to counter ethanol toxicity will be useful for bioengineering strains that can tolerate higher levels of ethanol.

Existing literature documents that organellar shape remodeling occurs in response to environmental stress. Such changes have been reported in Golgi (Acharya *et al*., 1998), peroxisomes (Kuravi *et al*., 2006), mitochondria (Bleazard *et al*., 1999), and lysosomes (Ward *et al*., 1997). Insights into such shape changes have also been reported for the vacuoles of *Saccharomyces cerevisiae* in response to external perturbations, such as oxidative stress (Corson *et al*., 1999), nutrient deprivation (Henne *et al*., 2015), thermal stress (Keuenhof *et al*., 2022), hypoosmotic shock, and salt stress (Li and Kane, 2009). This organelle is not only important for the turnover of various macromolecules but also for maintaining cellular pH and ion homeostasis (Li and Kane, 2009). Additionally, vacuole morphology is a fundamental part of this organism’s stress response and cell division. Under normal conditions, the vacuole forms two to three lobes and this multilobed morphology is the result of a dynamic equilibrium between the processes of vacuolar fission and fusion (Baars *et al*., 2007; Aufschnaiter and Büttner, 2019). Fission is important for vacuole inheritance as the vacuoles of the mother cell pinch off into small vesicles, which then migrate to the daughter cell, where they undergo homotypic vacuole fusion to form a new vacuole (Weisman, 2003). During hyperosmotic stress and oxidative stress, the vacuole fragments into several compartments, while it forms a single large compartment during hypoosmotic stress and starvation (Corson *et al*., 1999; Li and Kane, 2009; Henne *et al*., 2015). Thermal stress caused a >2x increase in the vacuolar size (Keuenhof *et al*., 2022).

Two derivatives of phosphatidylinositol (PtdIns), phosphatidylinositol 3-phosphate (PtdIns3P), and phosphatidylinositol 3,5-bisphosphate (PtdIns(3,5)P_2_) are important regulators of vacuole shape. PtdIns3P, produced by the lipid kinase Vps34, is present in the cytoplasmic leaflet of the endosomal and vacuolar membranes (Nascimbeni *et al*., 2017). PtdIns(3,5)P_2_ is enriched on the surfaces of late endosomes and the vacuole (Ho *et al*., 2012). It is generated from PtdIns3P by the phosphatidyl-inositol 3-phosphate 5-kinase, Fab1 (PIKFyve in mammals) (Gary *et al*., 1998; Zolov *et al*., 2012). This lipid kinase is part of the Fab1 complex comprising Fab1, the integral membrane protein Vac7, the scaffold protein Vac14, and the PtdIns(3,5)P_2_ phosphatase Fig4. Vac7, Vac14, and Fig4 are activators of Fab1 whereas Atg18, an effector of PtdIns(3,5)P_2_, is a negative regulator of the kinase maintaining a negative feedback loop to regulate levels of PtdIns(3,5)P_2_ (Efe *et al*., 2007; Botelho *et al*., 2008; Jin *et al*., 2008). Another negative regulator of Fab1 is the putative I-BAR domain-containing protein, Ivy1 (Malia *et al*., 2018). In addition to these extrinsic regulators, Fab1 activity is further regulated by intramolecular inhibitory interactions between its kinase and CCR domains (Lang *et al*., 2017). These intra- and inter-molecular associations precisely regulate the levels of PtdIns(3,5)P_2_ on the vacuoles. Under normal conditions, PtdIns(3,5)P_2_, a low abundance signaling lipid, constitutes only ∼0.05%-0.1% of total phosphoinositides present in the cell (Bonangelino *et al*., 2002; Duex *et al*., 2006). The levels of this lipid change dynamically to regulate the equilibrium between vacuole fusion and fission (Dove *et al*., 1997; Sbrissa *et al*., 1999; Bridges *et al*., 2012; Zolov *et al*., 2012). When vacuole fission is promoted during hyperosmotic stress, there is a substantial 10 to 20-fold elevation in the levels of PtdIns(3,5)P_2_, which causes vacuole fragmentation (Dove *et al*., 1997; Bonangelino *et al*., 2002). This fragmentation is essential to accommodate the loss in cell volume as water moves out of the cell. Fission is initiated by the formation of vacuole membrane invaginations that are enriched in PtdIns3P (Zieger and Mayer, 2012). Subsequently, PtdIns3P is converted to PtdIns(3,5)P_2_ by Fab1 (Gary *et al*., 1998). The PtdIns(3,5)P_2_ effector, Atg18, senses this lipid on the vacuole membrane and promotes membrane tubulation and scission (Gopaldass *et al*., 2017). During hypoosmotic stress, the low-abundance PtdIns(3,5)P_2_ is further suppressed, resulting in the downregulation of fission (Bonangelino *et al*., 2002). This shifts the dynamic equilibrium from fission towards fusion, leading to the formation of single-lobed vacuoles. Homotypic vacuole fusion, like any other membrane fusion event, requires pairing cognate SNAREs (soluble-N-ethylmaleimide-sensitive factor attachment protein receptor) proteins on the two vacuole compartments. After fusion, this *cis*-SNARE complex is uncoupled within the 20S complex, comprising the ATP-dependent NSF (*<u>N</u>*-ethylmaleimide-<u>s</u>ensitive <u>f</u>usion protein), its co-chaperonin α-SNAP, and the SNARE complex (Hohl *et al*., 1998; Ostrowicz *et al*., 2008).

Earlier reports have documented that ethanol exposure causes a transition from multilobed to single-lobed vacuoles in yeast (Meaden *et al*., 1999). We initiated this study to determine if this change is an actively managed process that is an integral part of the ethanol stress response of yeast cells. We observed that vacuole morphology changes very quickly in the presence of ethanol and occurs even when cells are exposed to low ethanol concentrations. This indicates that during ethanol stress, cells actively modulate the machinery regulating vacuole morphology. Vacuole enlargement is a part of ethanol stress response as conditional mutants with impaired vacuole fusion are more sensitive to ethanol. Furthermore, impaired vacuole fission contributes to the emergence of single-lobed vacuoles. This fission defect occurs due to a decrease in PtdIns(3,5)P_2_, brought about by modulation of the activity of Fab1. Fab1 activity is downregulated via the dissociation of the PtdIns(3,5)P_2_ effector Atg18 from the vacuolar membrane and inhibition in the recruitment of Atg18 onto the Fab1 complex.

## RESULTS

### Ethanol exposure results in enlarged vacuoles with invaginations

Wild-type yeast strains of opposite mating types, BY4741 and BY4742, exhibited sensitivity to 10% ethanol (Supplementary Figure 1). Therefore, we used this concentration of ethanol to monitor the change in vacuole morphology starting from 5 min post-ethanol addition, to 90 min. We stained yeast vacuoles of BY4742 using a lipophilic vital dye FM4-64 and observed the expected multilobed vacuole morphology (2-3 lobes) in the absence of ethanol (Figure 1A) (Vida and Emr, 1995; Aufschnaiter and Büttner, 2019). Even a brief exposure to ethanol for 5 min resulted in a change of vacuolar morphology from multilobed to a single sphere (Figure 1A). In addition to vacuole unlobing, ethanol exposure also caused a significant increase in the number of vacuoles where its limiting membrane invaginates into the lumen of the vacuole (Figure 1A). Three-dimensional image rendering and depth coding indicated that these membrane invaginations were deep, extending ∼2 µm from the vacuole surface into its lumen (Video 1 and 2). Image quantification revealed that compared to cells grown in the absence of ethanol, such invaginations of the vacuole membrane were present in a significant number of cells even after ethanol exposure for only 5 min, (Figure 1B). We observed a progressive increase in the number of cells with vacuolar membrane invaginations until 15 min of ethanol exposure, after which the number of cells with such membrane invaginations did not change substantially for up to 60 min. There was a drop in this number after 90 min (Figure 1B). We wanted to determine if the observed change in vacuole morphology with 10% ethanol also occurs when cells are exposed to lower concentrations of ethanol. Towards this, we monitored this organelle’s shape when cells were grown in ethanol concentrations ranging from 3% to 12% for 15 min (at this time point, 10% ethanol treatment caused the most invaginations). Exposure to even 3% ethanol resulted in a significant increase in the number of cells having the single-lobed phenotype rather than the multi-lobed morphology (Figure 1C). We also observed membrane invaginations of these single-lobed vacuoles (Figure 1D). The percentage of cells with single-lobed vacuoles increased with increasing ethanol concentration, up to 5%; thereafter, there was a decrease in the proportion of such cells as there was a rise in the proportion of cells with single-lobed vacuoles that displayed invaginations (Figure 1D). Thus, an increased number of vacuoles displayed membrane invaginations when cells were exposed to higher concentrations of ethanol. Taken together the data reveal that vacuole unlobing occurs very rapidly (<5 min), even at low ethanol concentrations (3%). In addition, invaginations of the unlobed vacuole membrane occur in an ethanol concentration-dependent manner, and the number of such invaginations increases with the time of ethanol exposure (up until 15 min) and then decreases.

**Figure 1.**
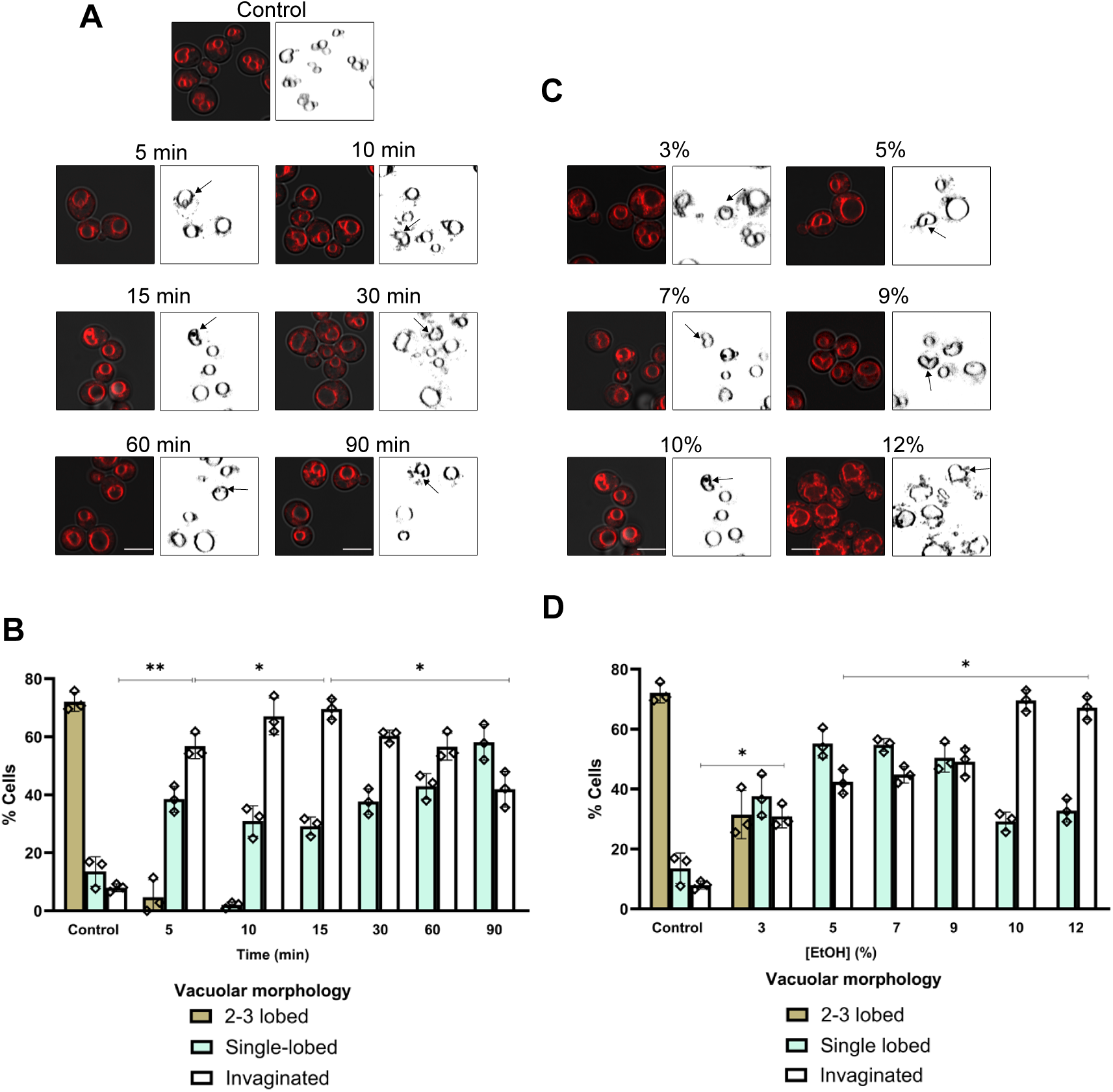
Ethanol stress response causes a change in the vacuolar morphology. (A) Vacuole membranes of BY4742 were stained with FM4-64 and vacuolar morphology was examined after treating the cells with 10% ethanol (v/v) at the indicated time points. The arrows indicate vacuolar invaginations. (B) Quantification of vacuole morphology as in A. (C) Vacuolar morphology of BY4742 was monitored under increasing concentrations of ethanol (3%-12%) for 15 min. The arrows indicate vacuolar invaginations. (D) Quantification of vacuole morphology as in C. In all cases, error bars represent s.d. of three individual experiments and each rhombus represents the mean of an independent biological replicate. **\***P<0.05; **\*\***P<0.01 (Paired two-tailed Student’s *t*-test). Scale bar: 5 µm.

### Vacuole shape change is essential for survival in the presence of ethanol

Since vacuole shape change, from multi-lobed to single-lobed, occurs within a short span of time, it is likely to involve homotypic vacuole fusion. Also, the appearance of an enlarged vacuole, even upon exposure to low levels of ethanol, indicates that this change in morphology may be an integral part of cellular response to ethanol. To explore if ethanol-induced vacuole fusion is a part of the ethanol stress response pathway, we determined if cells with defects in vacuole fusion are more sensitive to ethanol. We used two mutants, *sec18-1* and *sec17-1*, which encode temperature-sensitive versions of AAA-ATPase NSF and its co-factor α-SNAP, respectively. These associate with the *cis*-SNARE complex and utilize the energy from ATP hydrolysis to drive SNARE complex disassembly and the subsequent priming of the unpaired SNAREs (Ostrowicz *et al*., 2008). In the absence of this ATPase activity, vacuole fusion is hampered due to the limiting amounts of unpaired SNAREs. Both *sec18-1* and *sec17-1* are inviable at 37°C. At the permissive temperature of 24°C, the mutants behaved similarly to wild-type cells as their vacuole morphology was indistinguishable from the wild-type and exposure to 10% ethanol for 30 min caused their vacuoles to change from multilobed to a single-lobed compartment (Figure 2A, 2B, 2C, and 2D). Also, the growth of two of these two mutant strains was identical to that of the wild-type in the presence of 5%, 7%, and 9% ethanol (Figure 2E). At 26°C, while the growth of both mutants was similar to the wild-type in the absence of ethanol, they exhibited growth defects in its presence (Figure 2E). The *sec17-1* mutant was less sensitive than *sec18-1*, with growth defects evident only at 7% and 9%, while *sec18-1* showed clear growth retardation at 5% ethanol (Figure 2E). We observed a significant decrease in the number of *sec17-1* cells with single-lobbed vacuole after 30 min of exposure to 10% ethanol, from around 90% of cells exhibiting this phenotype at 24°C to around 50% at 26°C (Figure 2B). The reduction in cells capable of undergoing vacuole shape change upon ethanol exposure was greater for *sec18-1*, from ∼90% to <25% (Figure 2D). This indicates that even at a temperature permissive for growth (without ethanol), these mutants are sensitive to ethanol and have a defect in vacuole fusion; this fusion defect is more severe in the *sec18-1* mutant than in *sec17-1*. At 28°C, the growth of *sec17-1*, in the absence of ethanol, was indistinguishable from that of the wild-type, but *sec18-1* was unable to grow (Figure 2E). The negative impact of ethanol on the growth of *sec17-1* was evident even in the presence of 5% (Figure 2E). Both mutants exhibited fragmented vacuoles in the absence of ethanol, and there was no change in this morphology following ethanol exposure (Figure 2A to 2D).

**Figure 2.**
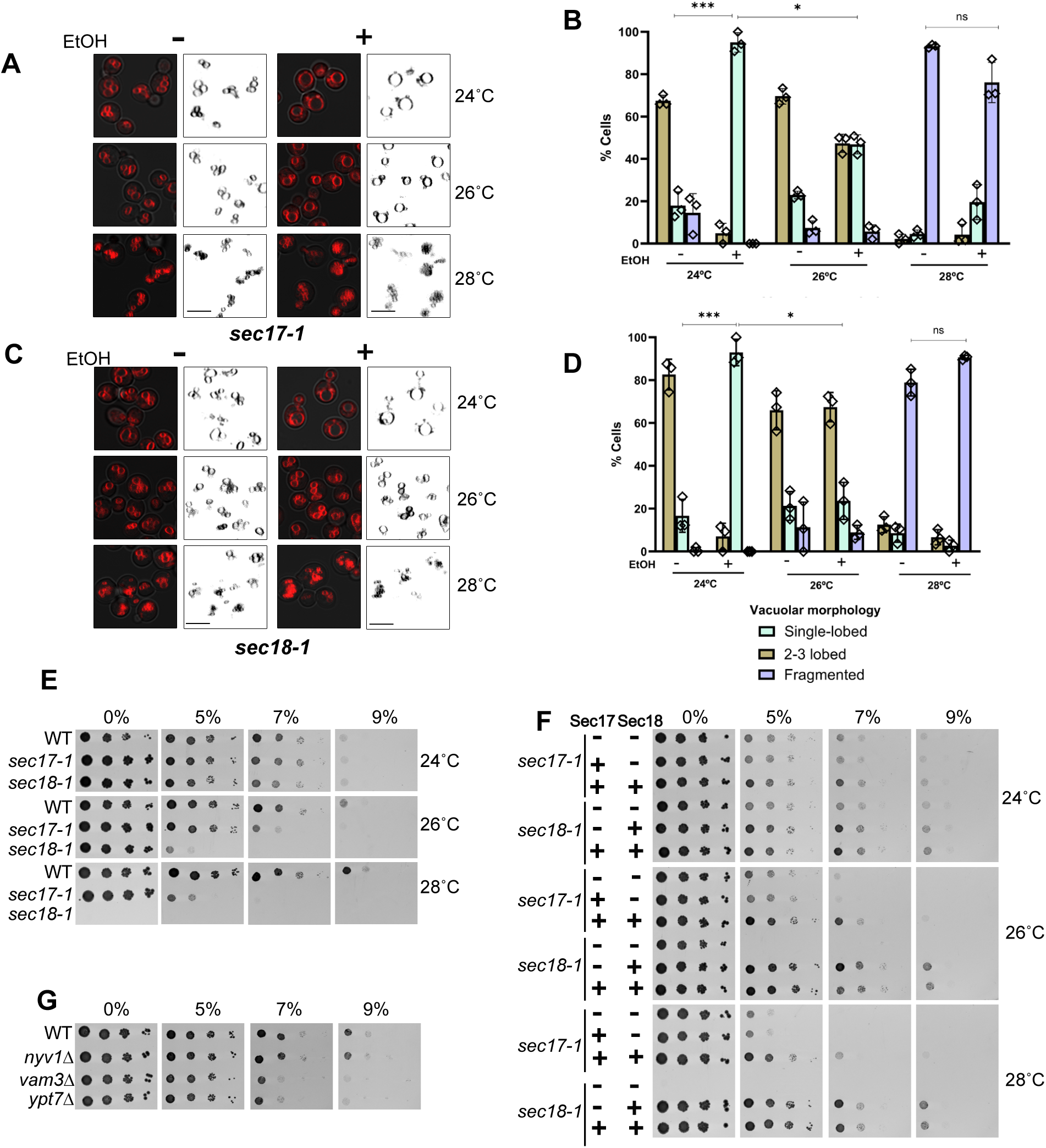
Vacuole enlargement is essential for cell survival in the presence of ethanol. (A) Vacuole membranes of *sec17-1* were stained with FM4-64 and vacuolar morphology was examined after treatment with 10% ethanol (v/v) for 30 min at three different temperatures (24°C, 26°C and 28°C). (B) Quantification of vacuole morphology as in A. (C) Vacuole membranes of *sec18-1* were stained with FM4-64 and vacuolar morphology was examined after treatment with 10% ethanol for 30 min at three different temperatures (24 °C, 26 °C and 28 °C). (D) Quantification of vacuole morphology as in C. (E) Growth of *sec17-1*, *sec18-1* and their isogenic wild-type. Cells were sequentially diluted 10x, spotted on synthetic complete medium containing varying concentrations of ethanol, and incubated at three different temperatures (24°C, 26°C and 28°C) for three days. (F) Growth rescue of temperature-sensitive *sec17-1* and *sec18-1* mutant strains. *sec17-1* and *sec18-1* mutants were co-transformed with constructs expressing Sec17, Sec18 or the empty vectors, which served as negative control. Cells were sequentially diluted 10x and spotted on synthetic minimal medium without leucine and uracil. The plates also contained varying concentrations of ethanol. The plates were incubated at three different temperatures (24°C, 26°C and 28°C). (G) Growth analysis for *nyv1*Δ, *vam3*Δ, and *ypt7*Δ. Cells were sequentially diluted 10x and were spotted on a synthetic complete medium containing varying concentrations of ethanol and were incubated at 28°C. In all cases, error bars represent s.d of three individual experiments and each rhombus represents the mean of an independent biological replicate. **\***P<0.05; **\*\***P<0.01; **\*\*\***P<0.001; **ns** no significant difference (Paired two-tailed Student’s *t*-test). Scale bar: 5 µm.

Both Sec17 and Sec18 have functions beyond that of membrane fusion. Given these diverse roles, the mutants are expected to be pleiotropic in nature (Haas, 1998; Zhao *et al*., 2007; Miao *et al*., 2013; de Paola *et al*., 2019). To establish that sensitivity to ethanol arises only due to vacuole fusion defect and not due to malfunctioning of other cellular events regulated by Sec17 or Sec18, we exploited previous observations that excess Sec17 blocks the docking of primed vacuoles and that Sec17 release from the vacuole membrane is driven by Sec18 early in the fusion reaction (Mayer *et al*., 1996; Wang *et al*., 2000; Baker and Hughson, 2016). Based on this, we hypothesize that overexpression of Sec17 in the *sec17-1* mutant will be unable to rescue the ethanol sensitivity as the excess pool of Sec17 will inhibit vacuole fusion, while the overexpression of Sec18 alone will rescue the ethanol sensitivity of *sec18-1* as these cells will not have any excess Sec17 and the endogenous pool of Sec17 will be sufficient to activate the wild-type Sec18 that has titrated out the mutant Sec18-1 from the 20S complex. Vacuole fusion arrest resulting from an excess of Sec17 can be overcome by overexpressing Sec18 (Wang *et al*., 2000). Hence, the ethanol sensitivity of *sec17-1* cells will be rescued when both Sec17 and Sec18 are overexpressed together. A similar overexpression of both proteins will not cause any difference in ethanol sensitivity when compared to the sensitivity of *sec18-1* cells overexpressing Sec18 alone. Consistent with this hypothesis, we observed that upon overexpression of Sec17 in the *sec17-1* mutant, there was no change in ethanol sensitivity at either 26°C or 28°C as the growth of these cells was comparable to that of *sec17-1* strain harboring only the vector (Figure 2F). However, overexpression of Sec18 in *sec18-1* caused a complete reversion of not only ethanol sensitivity (Figure 2F) but also temperature sensitivity as these cells could now survive at 28°C, a condition that was restrictive for the *sec18-1* mutant (Figure 2E). In keeping with our hypothesis, simultaneous overexpression of both Sec17 and Sec18 in the *sec17-1* mutant resulted in resistance to ethanol as these cells exhibited better growth (Figure 2F). Interestingly, we noted that *sec17-1* cells overexpressing the two proteins were more sensitive to ethanol than *sec18-1* cells overexpressing the two proteins; the growth of the latter was comparable to the wild-type. We are unable to account for this difference in growth of *sec17-1* and *sec18-1*.

As an independent confirmation of our hypothesis that vacuolar fusion is integral to ethanol stress response, we tested the ethanol sensitivity of other mutants with defects in this process. Following the activity of the 20S complex fusion-competent unpaired SNAREs are brought in closed proximity with the help of the Rab7-like GTPase Ypt7 to form the *trans*-SNARE complex, which then drives vacuole fusion by the formation of the *cis*-SNARE complex (Ostrowicz *et al*., 2008). We tested if deletion mutants lacking Nyv1 (v-SNARE), Vam3 (t-SNARE), or Ypt7 also exhibit enhanced sensitivity to ethanol. Except for *nyv1*Δ, all other mutants showed growth defects in the presence of ethanol when compared to wild-type (Figure 2G). The resistance of *nyv1*Δ was likely due to the presence of a bypass through Ykt6, which carries on vacuole fusion in the absence of Nyv1 (Dietrich *et al*., 2004; Watanabe *et al*., 2024). Consistently, unlike *vam3*Δ and *ypt7*Δ, the vacuoles of *nyv1*Δ cells enlarged following exposure to 10% ethanol for 30 min (Supplementary Figure S2). This is consistent with our interpretation that the enlarged vacuoles observed in ethanol-treated cells are a consequence of homotypic vacuole fusion and this is necessary for cells to withstand ethanol stress. Based on the above observations, we conclude that the enlargement of the vacuole upon ethanol exposure is an integral part of ethanol stress response as mutants that fail to unlobe their vacuole in the presence of ethanol exhibit enhanced ethanol sensitivity.

### Ethanol stress response involves inhibition of the vacuole fission machinery

Vacuole shape results from a dynamic equilibrium between the fission and fusion of vacuolar compartments. Ethanol stress causes a shift in this equilibrium toward unlobing, which may be due to the inhibition of vacuole fission, the promotion of vacuole fusion, or even both. To test if ethanol exposure has a negative effect on vacuole fission, we tested if fission-inducing agents can cause vacuole fission in ethanol-stressed cells. For induction of fission, we treated cells with 0.5 M NaCl, for 10 and 20 min, as such treatment is documented to promote rapid vacuole fission to counter osmotic imbalance (Michaillat *et al*., 2012). Consistent with previous observations, the presence of NaCl in the medium resulted in vacuole fragmentation in all the cells (Figure 3A). However when cells were first treated with ethanol for 5 min and then exposed to NaCl for 10 min, little or no fragmentation was observed; vacuoles were either multilobed or had single lobes, with or without invaginations (Figure 3A and 3B). This indicates that prior exposure to ethanol results in vacuoles that are compromised for fission. When the NaCl treatment was extended to 20 min, the number of cells having vacuoles with more than three lobes increased substantially to ∼40% (Figure 3B). Thus, the inhibition of vacuole fission upon exposure to 10% ethanol for 5 min appears reversible as it can be overcome by prolonging the exposure to NaCl. However, when the time of ethanol exposure was increased to 10 min, followed by either 10 or 20 min of salt shock, vacuolar fragmentation did not occur; instead, the unlobed vacuoles exhibited invagination (Figure 3A and 3B). Such vacuolar membrane invaginations are similar to those previously observed in various fission-deficient strains (Zieger and Mayer, 2012). Thus, exposure to ethanol for even 5 min inhibits vacuole fission, and this inhibition becomes more pronounced with an increase in the time of ethanol treatment. Based on the above observations, we conclude that ethanol-induced inhibition of the vacuole fission machinery contributes to the appearance of unlobed vacuoles. Since hypoosmotic shock also results in the unlobing of yeast vacuoles, we tested if there is a difference between water- and ethanol-induced vacuole unlobing. When we treated cells with water for 5 min, the change in their vacuole morphology was similar to that observed for ethanol exposure (compare Figure 3B and 3D). However, a comparison of the vacuole morphology of cells exposed to either water or ethanol for 10 min indicates that the latter treatment resulted in more invaginations compared to water-exposed cells. When water-treated cells were exposed to 0.5 M NaCl, for either 10 or 20 min, there was a substantial increase in the number of cells having fragmented vacuoles compared to the number of such cells after treatment with ethanol for 5 min (compare Figure 3B and 3D). Even after 10 min of hypoosmotic shock, followed by exposure to NaCl for 10 and 20 min, vacuole fragmentation was observed. In addition, only a minor population of cells exhibited vacuoles with invaginations indicating that, in comparison to ethanol, water-induced inhibition of vacuole fission is less severe and can be reversed more easily (Figure 3B and 3D). We also assessed the time taken by both ethanol- and water-treated cells to regain the normal multi-lobed vacuole shape by shifting to YPD for 10 and 20 min. Consistent with the NaCl treatment data, we observed that water-treated cells were able to regain multi-lobed vacuole morphology in the presence of YPD sooner than ethanol-treated cells (Supplementary Figure S3). Thermal stress also causes the unlobing of yeast vacuoles. We tested if cells exposed to thermal stress (37°C and 40°C) for 1 h retained the ability to undergo vacuole fragmentation following hyperosmotic stress with NaCl. Like ethanol stress, thermal stress also resulted in a block in vacuole fragmentation (Figure 3E and 3F). Thus, although vacuole unlobing is observed following various stress conditions, the underpinnings of the process are not the same.

**Figure 3.**
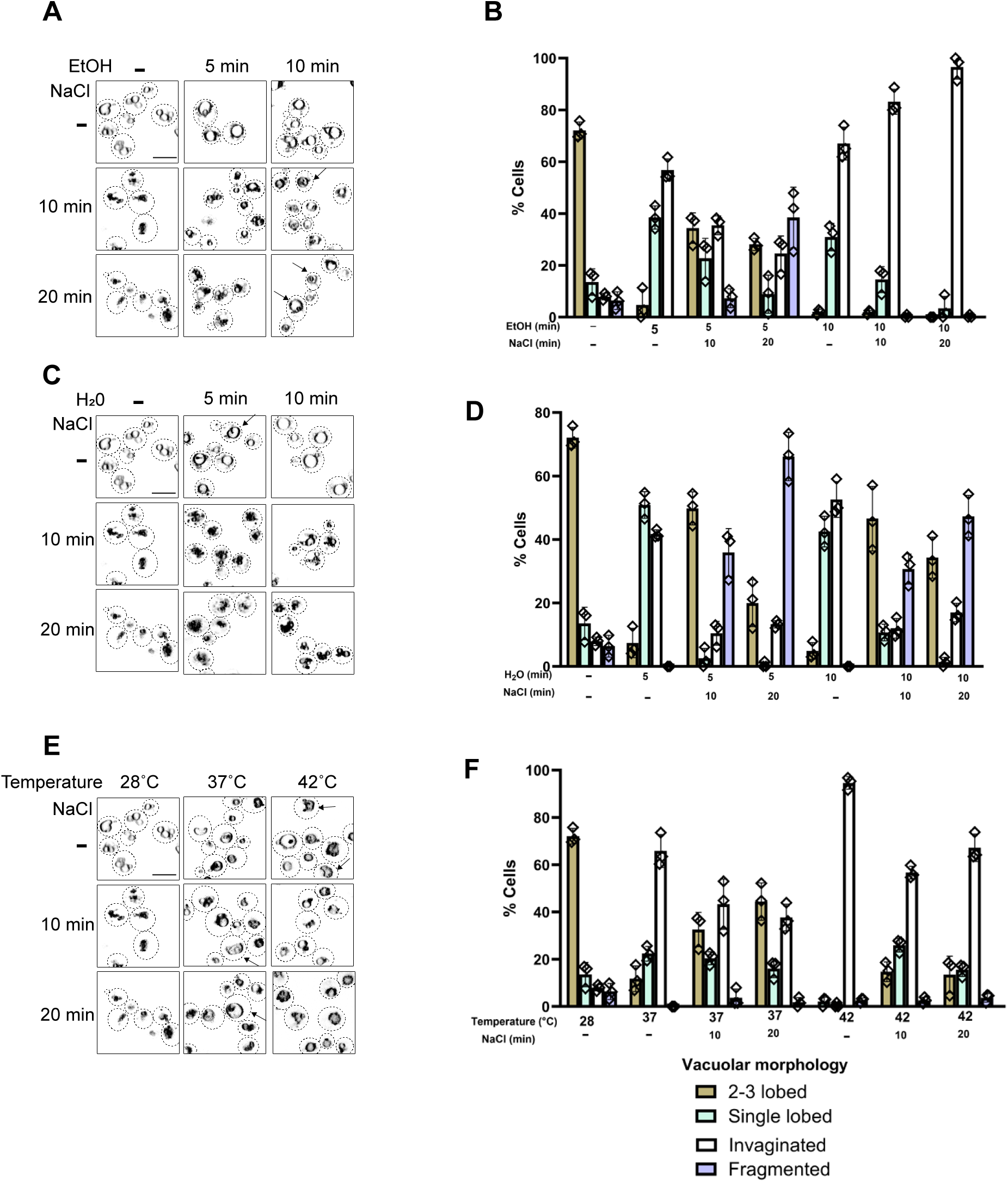
Ethanol inhibits the vacuole fission machinery. (A) Vacuole membranes of BY4742 were stained with FM4-64 and vacuole morphology was examined in cells treated with ethanol for 5 min and 10 min followed by treatment with 0.5 M NaCl for either 10 min or 20 min. The arrows indicate vacuolar invaginations. (B) Quantification of vacuole morphology as in A. (C) Vacuole morphology of cells exposed to hypoosmotic stress for 5 min and 10 min followed by treatment with 0.5 M NaCl for either 10 min or 20 min. (D) Quantification of vacuole morphology as in C. (E) Vacuole morphology of cells subjected to elevated temperatures of 37°C or 42°C for 1 h followed by treatment with 0.5 M NaCl for either 10 min or 20 min. (F) Quantification of vacuole morphology as in E. In all cases, error bars represent s.d. of three individual experiments and each rhombus represents the mean of an independent biological replicate. Scale bar: 5 µm.

### Ethanol stress response causes a redistribution of the phosphatidylinositol-3,5-bisphosphate sensor

Next, we wanted to understand how ethanol affects vacuole fission. Earlier studies have demonstrated that vacuolar fragmentation is initiated by the formation of membrane invaginations that are enriched for PtdIns3P. Next, the vacuole membrane protrusions that occur adjacent to these invaginations start to vesiculate into small structures, and this vesiculation depends on the conversion of PtdIns3P to PtdIns(3,5)P_2_ (Gary *et al*., 1998; Bonangelino *et al*., 2002). Given the role of these two lipids in vacuole fragmentation, we wanted to determine if ethanol-induced fission block results in any perturbations in the distribution of these two lipid regulators.

To determine the distribution of PtdIns3P, we used a PtdIns3P-binding sensor having two copies of the FYVE domain of the human Hrs protein fused to GFP (FYVE_2_-GFP) (Zieger and Mayer, 2012). Consistent with previous reports, we observed that FYVE_2_-GFP localized to both endosomes and the vacuole membrane of wild-type cells, while no membrane-associated GFP signal was evident in *vps34*Δ cells where no PtdIns3P was produced (Figure 4A). Upon exposure to 10% ethanol for 30 min, the GFP signal was associated with the membranes of unlobed vacuoles and also in the membranes lining the deep invaginations of the vacuole membrane (Figure 4A). Thus, ethanol stress does not affect the production of PtdIns3P or its distribution at the vacuolar membrane. Such distribution of PtdIns3P to the vacuolar membrane and the membrane invaginations was also observed in *fab1*Δ cells, both with and without ethanol stress (Figure 4A). This is consistent with previous reports that *fab1*Δ cells, with negligible levels of PtdIns(3,5)P_2_, also form these membrane invaginations under normal conditions. However, in their case, these invaginations persist, and these deformations do not bud off to form vacuolar fragments (Zieger and Mayer, 2012).

**Figure 4.**
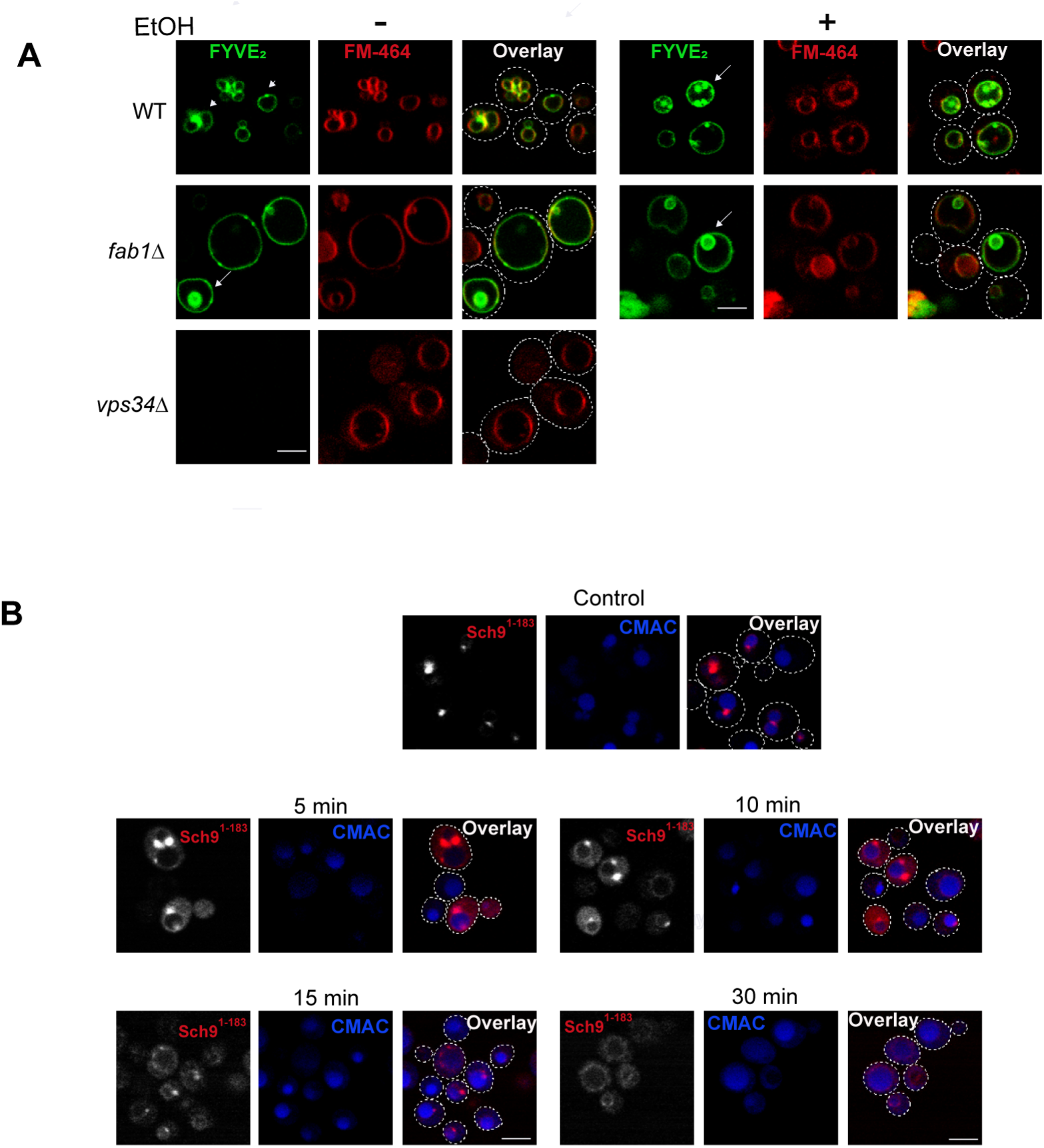
Ethanol causes the redistribution of the phosphatidylinositol-3,5-bisphosphate sensor. (A) Subcellular distribution of PtdIns3P after ethanol stress. FYVE_2_-GFP was expressed in wild-type, *fab1*Δ, and *vps34*Δ. Vacuole membranes were stained with FM4-64 and cells were imaged after adding 10% ethanol (v/v) for 30 min. The arrows mark vacuolar invaginations and arrowheads mark endosomes. (B) Subcellular distribution of PtdIns(3,5)P_2_ after ethanol stress. Sch9^1-183^ fragment was C-terminally tagged with RFP and was expressed under a constitutive promoter in wild-type cells. Vacuolar lumen was stained with CMAC and cells were visualized after adding 10% ethanol (v/v) for indicated time points. Scale bar: 5 µm

Our earlier observation led to the conclusion that ethanol-treated cells have unlobed vacuoles with invaginations (Figure 1B), and these membrane deformations persisted even after 20 min of salt shock (Figure 3A), indicating a defect in vacuole fragmentation. This lack of fragmentation led us to hypothesize that ethanol stress modulates PtdIns(3,5)P_2_ pools on the vacuole membrane. We monitored the distribution of PtdIns(3,5)P_2_ on the vacuole membrane using a PtdIns(3,5)P_2_ sensor in which the N-terminal 183 residues of Sch9, a homolog of mammalian S6 kinase, is C-terminally tagged with RFP (Sch9^1-183^-RFP). This sensor localizes to perivacuolar dots in a PtdIns(3,5)P_2_-dependent manner as this signal was absent in *fab1*Δ, *vac7*Δ, *vac14*Δ, and *fig4*Δ, all of which have negligible PtdIns(3,5)P_2_ (Jin *et al*., 2014; Chen *et al*., 2021). Consistently, we also observed that these deletion strains lack Sch9^1-183^-RFP positive perivacuolar dots (Supplementary Figure S4). Upon ethanol treatment, this perivacuolar localization of Sch9^1-183^-RFP was affected (Figure 4B). After 5 and 10 min of ethanol addition, besides perivacuolar dots, we observed a cytoplasmic RFP signal (Figure 4B). 15 min after treatment, we observed only a few perivacuolar dots with low signal intensity. Instead, the signal was predominantly on the vacuole membrane and in the cytoplasm. By 30 min, there were almost no perivacuolar dots while the signal at the vacuole membrane and cytoplasm persisted (Figure 4B). This time-dependent progressive shift of Sch9^1-183^-RFP signal from perivacuolar dots to the vacuole membrane and the cytoplasm indicates that ethanol stress response involves a modulation of PtdIns(3,5)P_2_. Thus, while there was little or no change in the distribution of PtdIns3P in cells exposed to ethanol, there was a substantial change in PtdIns(3,5)P_2_ distribution.

### Ethanol-induced fission block is impaired in mutants with elevated levels of PtdIns(3,5)P_2_

To determine whether altered PtdIns(3,5)P_2_ distribution causes vacuole enlargement during ethanol stress, we tested if enhancing the pools of this lipid could prevent the formation of enlarged vacuoles post-ethanol stress. For this, we used mutants with high levels of this lipid due to dysregulation of the PtdIns(3,5)P_2_ synthesis machinery. Ivy1 is a known negative regulator of Fab1 (Malia *et al*., 2018). As *ivy1*Δ mutants have high levels of PtdIns(3,5)P_2_, under normal conditions these cells contain fragmented vacuoles instead of the two to three lobes observed in the wild-type (compare Figure 1A and Figure 5A) (Malia *et al*., 2018). Upon ethanol treatment, the vacuoles of *ivy1*Δ cells remained fragmented even after 15 min of ethanol treatment (Figure 5A). In contrast, wild-type cells exhibit enlarged vacuoles (with or without invaginations) after 5 min of ethanol treatment (compare Figure 1A and 5A). Only after 25 min of ethanol treatment, we observed vacuole enlargement in *ivy1*Δ (Figure 5A and 5B). This delay in vacuole enlargement is likely due to the enhanced PtdIns(3,5)P_2_ levels in this mutant that promote vacuole fission. Elevated levels of this lipid also allow *ivy1*Δ cells to undergo rapid vacuole fragmentation after 10 min of treatment with 0.5 M NaCl, whereas wild-type cells failed to undergo fragmentation even after 20 min (compare Figure 3A and Supplementary Figure S5). Consistently, the unlobed vacuoles of *ivy1*Δ cells regained their fragmented vacuole phenotype after shifting YPD for only 10 min, in contrast to wild-type cells, which even after 20 min of similar shift to YPD, did not regain their normal multilobed vacuole morphology (compare Supplementary Figure S3 and S5). These observations indicate that there is a delay in ethanol-induced block in vacuole fission in mutants with elevated pools of PtdIns(3,5)P_2_.

**Figure 5.**
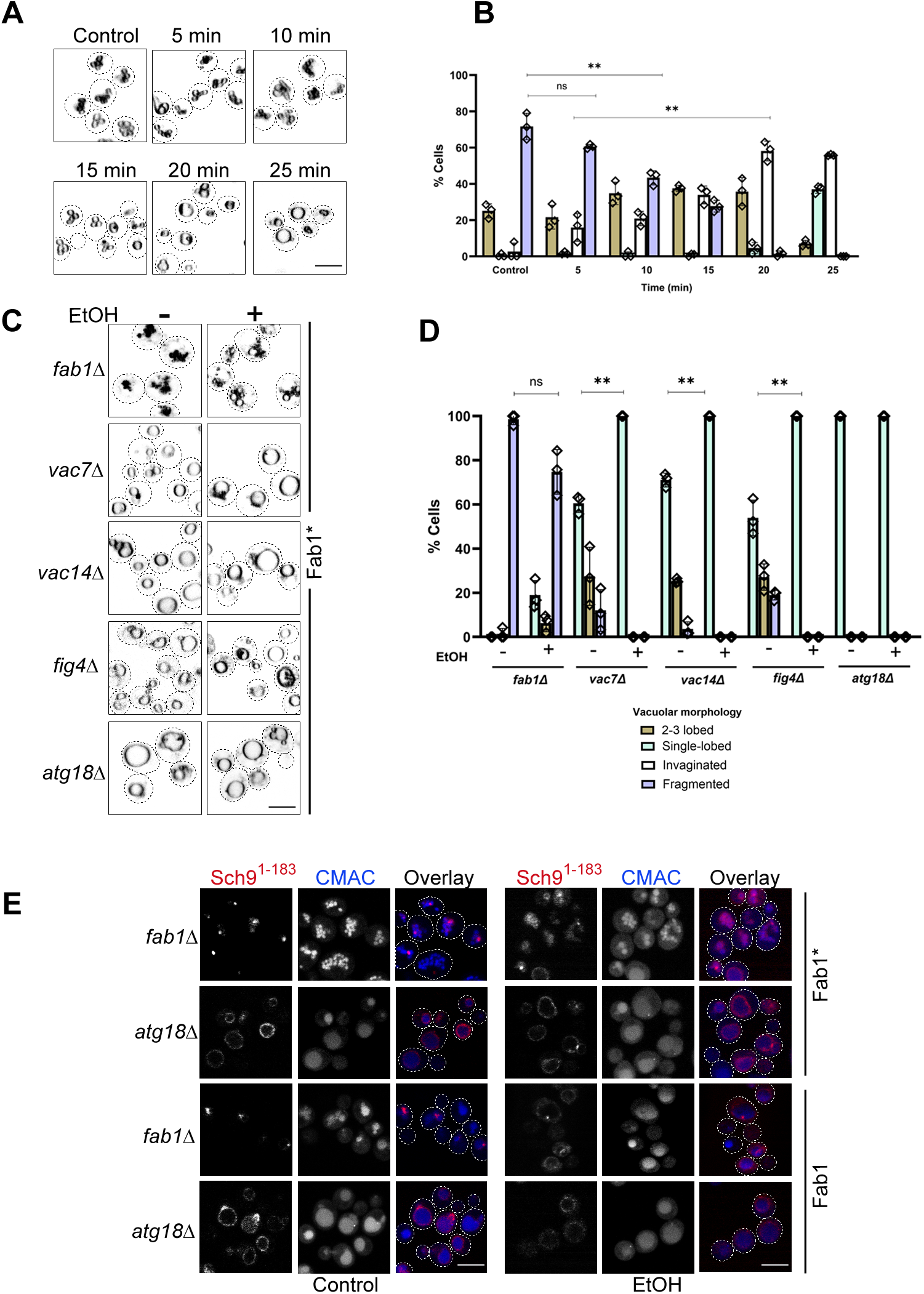
Elevated pools of PtdIns(3,5)P_2_ delay ethanol-induced fission block. (A) Vacuole membranes of *ivy1*Δ were stained with FM-464, and vacuole morphology of ethanol-treated ivy1Δ cells was examined at 10% ethanol (v/v) concentration, at the indicated time points. (B) Quantification of vacuole morphology as in A. (C) Vacuole morphology was examined for the individual deletion mutants of the Fab1 complex that were expressing the Fab1 hyperactive allele (Fab1*). (D) Quantification of vacuole morphology as in C. (E) Distribution of PtdIns(3,5)P_2_ as determined using Sch9^1-183^-RFP localization in *fab1*Δ and *atg18*Δ expressing either Fab1* or Fab1 in the presence or absence of ethanol. In all cases, error bars represent the s.d. of three individual experiments, and each rhombus represents the mean of an independent biological replicate. **\*\***P<0.01; **ns** no significant difference (Paired two-tailed Student’s *t*-test). Scale bar: 5 µm

To independently validate our previous data that mutants with elevated PtdIns(3,5)P_2_ show delay in ethanol-induced vacuole enlargement, we monitored the vacuole morphology in cells expressing a hyperactive mutant of Fab1 (Fab1*), which produces elevated levels of PtdIns(3,5)P_2_. Consequently, the expression of Fab1* is known to cause vacuole fragmentation (Gary *et al*., 2002; Malia *et al*., 2018). We analyzed the vacuolar morphology of various Fab1 complex mutants expressing Fab1*, in the presence or absence of ethanol. Consistent with previous literature, we observed vacuolar fragmentation in *fab1*Δ expressing Fab1* (Figure 5C). When these cells were treated with 10% ethanol for 30 min, the fragmented vacuoles enlarged marginally indicating that elevated levels of PtdIns(3,5)P_2_ hinder vacuole unlobing upon ethanol exposure (Figure 5C and 5D). However, this marginal enlargement indicates that there could still be some partial depression of even these elevated PtdIns(3,5)P_2_ levels in response to ethanol. Consistently, we observed perivacuolar distribution of Sch9^1-183^ even after 30 min of ethanol exposure whereas no such distribution was observed in wild-type cells (compare Figure 4B and Figure 5E, right panel). It may be noted that compared to no ethanol treatment control, there is some redistribution of Sch9^1-183^ in *fab1*Δ expressing Fab1* (Figure 5E, compare left and right panels), which again supports the notion that there is some reduction of the elevated PtdIns(3,5)P_2_ levels. This distribution pattern for Sch9^1-183^ was dependent on the dosage of PtdIns(3,5)P_2_ as vacuole enlargement, in the presence of ethanol was observed in *fab1*Δ cells expressing the wild-type Fab1 from a centromeric vector, which do not have elevated levels of PtdIns(3,5)P_2_ (Figure 5E). Such cells also had enhanced levels of Sch9^1-183^ in their cytoplasm and vacuole membrane, which is similar to the wild-type cells under comparable conditions (compare Figure 4B and 5E). These results indicate that elevated PtdIns(3,5)P_2_ pools facilitate the redistribution of the lipid sensor back to the perivacuolar dots from the cytoplasm even under ethanol stress.

Besides Ivy1, the activity of Fab1 kinase is also controlled by the other activators of the Fab1 complex, *viz.* Vac7 and Vac14, the lipid phosphatase Fig4 (converts PtdIns(3,5)P_2_ to PtdIns3P), and the PtdIns(3,5)P_2_ effector, Atg18. These proteins reside on the vacuole membrane and function together to tightly regulate PtdIns(3,5)P2 levels and, consequently, vacuole morphology (Botelho *et al*., 2008; Jin *et al*., 2008). *vac7*Δ, *vac14*Δ, and *fig4*Δ mutants contain negligible levels of PtdIns(3,5)P_2_ as the activation of Fab1 is hampered in the absence of these proteins (Rudge *et al*., 2004). In line with this, the expression of Fab1 in these three deletion strains (*vac7*Δ, *vac14*Δ, and *fig4*Δ) could not suppress the enlarged vacuole phenotype, whereas Fab1* could partially do so (Figure 5C and Supplementary Figure S6). Image quantification data indicated very little or no fragmentation as a majority of the cells had single-lobed vacuoles with only a minor population (∼30%) carrying 2-3 lobed vacuoles (Figure 5D). Exposing the cells to 10% ethanol for 30 min resulted in all cells having single-lobed vacuoles (Figure 5C and 5D). These data suggest that vacuolar enlargement under ethanol stress could not be suppressed by boosting PtdIns(3,5)P_2_ levels in mutants where the activation of the Fab1 complex was hampered.

In *atg18*Δ, we observed that the expression of either Fab1* or Fab1 resulted in enlarged vacuoles either in the presence or absence of ethanol (Figure 5C, 5D, and Supplementary Figure S6). Consistently, *atg18*Δ cells coexpressing either Fab1 or Fab1* and Sch9^1-183^-RFP exhibited signals on the vacuole membrane both with and without ethanol (Figure 5E). This was consistent with earlier reports documenting that Atg18 is required to shift this sensor from the vacuole membrane to the perivacuolar dots (Chen *et al*., 2021). Also, Atg18 is a known effector of PtdIns(3,5)P_2_, responsible for membrane septation (Gopaldass *et al*., 2017). *atg18*Δ cells have been shown to contain high levels of PtdIns(3,5)P_2_ due to the impaired turnover of this lipid (Efe *et al*., 2007). Elevating PtdIns(3,5)P_2_ further by expressing Fab1* does not cause any change in the distribution of Sch9^1-183^-RFP under ethanol stress. This is because, in the case of *atg18*Δ, ethanol cannot cause any reduction of these double-enhanced PtdIns(3,5)P_2_. This is in contrast to *fab1*Δ expressing Fab1*, where there was a minor redistribution of the sensor due to some reduction of the elevated PtdIns(3,5)P_2_ levels upon ethanol treatment (Figure 5E). However, this supports our observation of little or no alteration in the distribution of the Sch9^1-183^-RFP sensor in *atg18*Δ expressing either Fab1 or Fab1* even after the addition of ethanol (Figure 5E). Thus, elevating the levels of the ligand PtdIns(3,5)P_2_ can prevent vacuole enlargement during ethanol stress provided the effector, Atg18, is present. In its absence, vacuole enlargement occurs even when PtdIns(3,5)P_2_ levels are high. Taken together, these observations prove that vacuole enlargement during ethanol stress response involves suppression of PtdIns(3,5)P_2_ levels.

### Modulation of phosphatidylinositol-3,5-bisphosphate during ethanol stress response is via the dissociation of Atg18

The regulated production of PtdIns(3,5)P_2_, requires the physical association of Vac14, Vac7, Fig4, and Fab1, together with Atg18 and other regulators such as Ivy1. The scaffold adaptor protein Vac14 nucleates this Fab1 complex assembly (Jin *et al*., 2008; Alghamdi *et al*., 2013). To understand how cells modulate levels of PtdIns(3,5)P_2_ during ethanol stress, we checked the assembly of the regulators of Fab1 onto the Vac14 scaffold. For this, we have used the yeast two-hybrid assay. This assay was initially used to establish the interactions between the Fab1 complex components and also the interactions between the various domains of Fab1 (Jin *et al*., 2008; Lang *et al*., 2017). The proteins were expressed as fusions of either the Gal4-activation domain (AD) or the Gal4-DNA binding domain (BD). Transformants expressing various binary combinations of these fusions were assessed for expression of the two reporter genes, *HIS3* and *ADE2*, in the presence or absence of 5% ethanol. Consistent with previous reports, we observed the interaction of Vac14 with Vac7, Fig4, and Atg18, and also with itself (Figure 6A). We did not observe any interaction between Vac14 and Ivy1. The interaction between Vac14 and Vac7/Fig4/Atg18/Vac14 was strong as all the transformants expressed not only the *HIS3* reporter but also the *ADE2* reporter; the latter is expressed only when the two interacting proteins have a strong affinity for each other. When the same assay was performed in the presence of 5% ethanol, there was no change in the interaction of Vac14 with either itself or Vac7 or Fig4 as growth was observed in the absence of either histidine or adenine (Figure 6A). However, the interaction with Atg18 was completely abolished as these transformants did not express even the *HIS3* reporter (Figure 6A). It may be noted that the interaction between Atg18 and Vac14 detected via yeast two-hybrid assay is independent of Atg18’s interaction with its ligand PtdIns(3,5)P_2_ as this interaction occurs within the confines of the nucleus and not in the context of the vacuole membrane surface. This indicates that during ethanol stress, cells may modulate the levels of PtdIns(3,5)P_2_ by preventing the recruitment of Atg18 to the Fab1 complex, and this mode of modulation is independent of Atg18 binding to PtdIns(3,5)P_2_.

**Figure 6.**
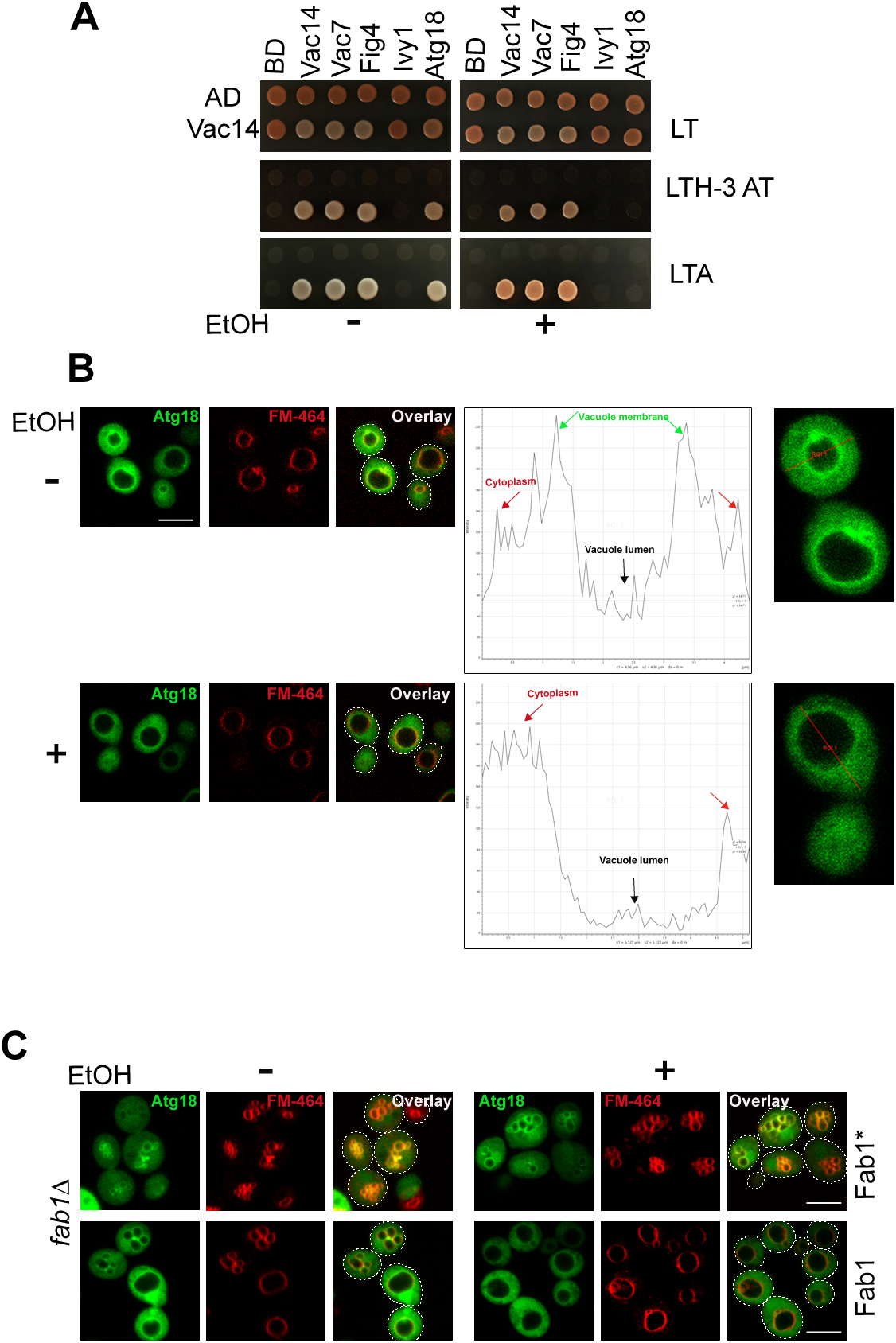
Ethanol stress response entails an inhibition of Atg18 recruitment onto the Fab1 complex. (A) Yeast-two hybrid assay for the interaction of Vac14 fused with the Gal4-activation domain (AD) with itself or Vac7, Vac14, Fig4, and Atg18 that were fused with the Gal4-DNA binding domain (BD). Binary combinations of the indicated AD and BD fusion proteins were coexpressed in PJ69-4A cells. The interaction between various binary fusion combinations was assessed by the expression of the *HIS3* and *ADE2* reporter genes by monitoring the growth of the transformants on plates lacking leucine, tryptophan, and histidine with 2.5 mM 3-AT (LTH^—^3-AT), or lacking leucine, tryptophan, and adenine (LTA^-^), in the presence and absence of ethanol respectively. (B) Localization of Atg18-GFP expressed under a galactose inducible promoter was monitored following treatment with 10% ethanol for 30 min. Lines indicate the region of interest (ROI) for the intensity plots for a particular *z*-stack image. The arrows mark the intensities of signals for the cytoplasm, vacuole membrane, and vacuole lumen. (C) Localization of Atg18-GFP in *fab1*Δ expressing either Fab1* or Fab1 in the presence or absence of ethanol.

If there is a lack of interaction between Atg18 and Vac14, then it is likely that Atg18 will not be recruited to the Fab1 complex in ethanol-treated cells. To check if the subcellular distribution of Atg18 is affected upon exposure to ethanol, we monitored the distribution of Atg18-GFP in the presence and absence of ethanol. Consistent with previous reports, we observed that under normal conditions, Atg18-GFP was present in the cytoplasm and also at the vacuole membrane (Figure 6B) (Efe *et al*., 2007). There was a change in the distribution when cells were exposed to 10% ethanol for 30 min. While the cytoplasmic pool persisted, that at the vacuole membrane was lost. The intensity plots also show that ethanol exposure results in the loss of Atg18-GFP signal from the vacuole membrane (Figure 6B). This indicates that ethanol stress modulates endogenous PtdIns(3,5)P_2_ pools by causing the dissociation of Atg18 from the vacuole membrane. This observation is consistent with our yeast two-hybrid assay results that show selective inhibition of the interaction between Vac14 and Atg18 in the presence of ethanol.

To test if ethanol-induced alteration in the pools of this lipid plays a role in the relocalization of Atg18 during ethanol exposure, we determined the distribution of Atg18-GFP in cells expressing Fab1*, in the presence and absence of ethanol. We observed that unlike in wild-type cells, where the vacuole membrane signal for Atg18-GFP was lost following ethanol treatment, in cells expressing Fab1*, the vacuole membrane signal for Atg18-GFP remained unchanged with or without ethanol (Figure 6C). This proves that augmenting recruitment of Atg18 to the vacuole limiting membrane via enhanced levels of PtdIns(3,5)P_2_ caused vacuole fragmentation even under ethanol stress (Figure 6C). Consequently, this membrane recruitment of Atg18-GFP was dependent upon the levels of PtdIns(3,5)P_2_ since expression of Fab1, instead of Fab1*, did not result in the persistence of the vacuole membrane signal after ethanol exposure, which is similar to wild-type cells (Figure 6B and 6C). We observed a similar change in the distribution of the PtdIns(3,5)P_2_ sensor, Sch9^1-183^-RFP expressing Fab1* or Fab1 under ethanol stress (Figure 5E). Thus, the unlobing of the vacuole in response to ethanol stress involves a block in vacuole fission and this block is achieved through the combined effect of reduction in PtdIns(3,5)P_2_ levels and the loss of interaction between Vac14 and Atg18.

## DISCUSSION

Previous studies have documented that vacuoles enlarge upon exposure to ethanol (Meaden *et al*., 1999). This study has shown that the change in the morphology of the vacuole is very rapid and occurs even in the presence of low concentrations of ethanol. Also, mutants defective for homotypic vacuole fusion are more sensitive to ethanol. These observations lead us to conclude that change in vacuole morphology is an integral part of cellular response to ethanol toxicity. Cells execute this rapid change by inhibiting the vacuole fission machinery by suppressing PtdIns(3,5)P_2_ levels, which hampers vacuole membrane recruitment of Atg18. Membrane recruitment of Atg18 is further reduced as its interaction with the Vac14 scaffold protein is hampered in the presence of ethanol.

Why would the vacuole need to undergo such enlargement in the presence of ethanol? Our data shows that cells enlarge their vacuoles via an active process in response to ethanol and this can only happen if such a change in shape helps the cell counter deleterious change(s) that occur in ethanol’s presence. An example of such an ethanol-induced deleterious change is alterations in the physical properties of cellular membranes. The hydrophobic nature of ethanol increases membrane fluidity (Goldstein, 1986; Tóth *et al*., 2014; Navarro-Tapia *et al*., 2018). Fluidity decreases membrane tension, which is an important parameter for regulating the activities of various membrane-spanning transporters and channel proteins (Chang *et al*., 2010). The activities of such transporters and channels of the vacuole membrane are crucial during ethanol exposure as they play vital roles in maintaining homeostasis. A way to reestablish optimum vacuole membrane tension following ethanol exposure is through a decrease in vacuolar surface area-to-volume ratio. The formation of a single large compartment from the multilobed vacuole will result in a lowering of this ratio and a consequent increase in membrane tension. Thus, the increase in volume relative to the surface may be an early response aimed toward reestablishing the requisite membrane tension to ensure the continued functions of the membrane-associated proteins. While such a decrease in surface area to volume ratio may be a rapid response towards reestablishing tension, previous studies indicate that yeast cells make additional attempts to correct ethanol-induced fall in membrane tension. There is a positive correlation between the tension of biological membranes and the area occupied by the individual lipids within the membrane, i.e. area per lipid (APL) (Jalali *et al*., 2024). It has been previously reported that yeast cells increase levels of unsaturated fatty acids and ergosterol during ethanol stress (Ding *et al*., 2009). Since unsaturated fatty acids have higher APL, the presence of these in membranes will result in higher tension (Lira *et al*., 2023). We propose that the increase in volume relative to the surface is likely to be a very early response aimed toward reestablishing the requisite membrane tension to ensure the continued functions of the membrane-associated proteins. This may be accompanied by an additional increasing unsaturated fatty acid content that will further augment membrane tension. It is worth noting that increasing tension via an increase in vacuolar volume has to be within the limits imposed by the volume restrictions for the entire cell. Incidentally, two other stress conditions, thermal and hypoosmotic stress, which increase membrane fluidity are also known to cause enlargement of vacuoles (Los and Murata, 2004). However, the mechanism by which this enlargement is executed is unlikely to be the same. Our results indicate that the block in vacuole fission following hypoosmotic stress is less severe than that caused by ethanol exposure whereas the block imposed by thermal stress is more detrimental (Figure 3). It may also be a reflection of the fact that ethanol or thermal stress may cause perturbations of a larger number of homeostatic parameters than hypoosmotic stress. Hence, it takes longer for all of these parameters to return to normal.

In addition to ethanol-induced vacuole enlargement, we also observed vacuoles that exhibited deep, tubular invaginations of the vacuole membrane into the lumen. As previously mentioned, such invaginations have been previously documented to precede vacuolar fragmentation events dependent on PtdIns(3,5)P_2_ and form very rapidly when cells encounter salt stress (Zieger and Mayer, 2012). In line with this, various fission-deficient strains with low or negligible levels of PtdIns(3,5)P_2_ appear to form these membrane invaginations under salt stress. However, in their case, these invaginations persist for a longer time than wild-type cells and these deformations do not bud off to form vacuolar fragments (Zieger and Mayer, 2012). Our observations indicate similar vacuolar invaginations even upon brief exposure to ethanol which implies that whether the invaginations are from fission-deficient strains or from ethanol-stressed cells whose fission has been blocked leading to a decrease in the PtdIns(3,5)P_2_ levels, the nature of invaginations are similar, proving that ethanol causes inhibition in vacuole fission (Figure 1). Consistently, the number of vacuoles with invaginations in ethanol-exposed cells subjected to subsequent salt shock showed a positive correlation with the time of ethanol exposure (Figure 3). Longer treatment with ethanol heightened membrane protrusions (80-100% of cells show invaginations) whereas brief exposure leads to a decrease in invaginations and an increase in vacuole fragments (Figure 3B). This indicates that ethanol-treated cells upon salt shock attempt vacuole fragmentation but are unable to complete the process due to the fission block. We also observed a PtdIns(3,5)P_2_-dependent alteration of Sch9 distribution from perivacuolar dots to the vacuole membrane and cytosol. The shift occurred over a period of 15 min (Figure 4B). This gradual shift is consistent with the positive correlation between ethanol treatment duration and YPD-mediated recovery (Supplementary Figure S3). Although the shift in Sch9 distribution is gradual, the system is likely to be sensitive such that a slight decrease in PtdIns(3,5)P_2_ can inhibit fission.

Time kinetics experiment revealed that there was a shift in the ratio of vacuoles having invaginations and those without, as the latter population increased with time relative to the former (Figure 1A). We speculate that a gradual decrease in PtdIns(3,5)P_2_ results in cells initially trying to fragment vacuoles as invaginations are maximal at 15 min but upon prolonged treatment, the invaginations decrease. Furthermore, ethanol stress causes enhanced cytoplasmic and vacuolar acidification defects (Charoenbhakdi *et al*., 2016). Previous reports document that the initiation of invaginations requires V-ATPase activity and it is dependent on PtdIns(3,5)P_2_ (Farge and Devaux, 1992; Sackmann and Feder, 1995; Li *et al*., 2014; Banerjee *et al*., 2019). Since ethanol stress response depletes PtdIns(3,5)P_2_, the assembly of the V_0_ and V_1_ subunits of V-ATPase is likely to be hampered. We hypothesize that at the initial exposure to ethanol, even though PtdIns(3,5)P_2_ pools have been reduced by downregulating its production, the amount of active V-ATPase is sufficient to initiate vacuole membrane invaginations. However, with prolonged exposure, the drop in the levels of PtdIns(3,5)P_2_ results in inactivation of V-ATPase, thus preventing the formation of new invaginations. Hence there is a rise in the population of cells carrying enlarged vacuoles without invaginations (Figure 1A). The development of vacuolar invaginations upon ethanol stress is also due to the modulation in the distribution of vacuolar lipids, which causes them to phase separate into two liquid phases (Leveille *et al*., 2022; Kim and Budin, 2024). It is thus conceivable that yeast cells actively remodel their membranes and undergo phase separation to adjust to environmental stresses.

The conversion of PtdIns3P to PtdIns(3,5)P_2_ is necessary for the vesiculation of invaginated vacuoles given if these lipids recruit other lipid-binding proteins onto the vacuole membrane that induces further shape change. One such lipid-binding protein is Atg18 which can bind to both PtdIns3P and PtdIns(3,5)P_2_ and can also regulate Fab1 activity (Dove *et al*., 2004). Atg18 has dual physiological functions, i.e., one is the regulation of autophagy via its interaction with Atg2 mediated by PtdIns3P and the second is the regulation of vacuole shape mediated by PtdIns(3,5)P_2_ (Watanabe *et al*., 2012; Rieter *et al*., 2013; Kotani *et al*., 2018). Atg18 can bind to these lipids via a seven-bladed ß-propeller motif (Dove *et al*., 2004). It has unique binding sites for the lipids on blades 5 and 6 and can embed in the vacuole membrane via the hydrophobic patches on blades 6 or 7 (Krick *et al*., 2006; Baskaran *et al*., 2012; Watanabe *et al*., 2012). Our results show that Atg18 membrane recruitment is hampered upon ethanol stress in a PtdIns(3,5)P_2_-dependent manner (Figure 6B). We have also shown that tethering Atg18 to the vacuole limiting membrane by elevating PtdIns(3,5)P_2_ promoted vacuole fragmentation during ethanol stress (Figure 6C). In addition, the binding of Atg18 to PtdIns(3,5)P_2_ also depends on its phosphorylation status as dephosphorylation of Atg18 promotes binding to this lipid thereby causing vacuole fission (Tamura *et al*., 2013). Our results have uncovered that the recruitment of Atg18 to the vacuole membrane may also be modulated via its binding to Vac14 (Figure 6A). This interaction is PtdIns(3,5)P_2_-independent, therefore it may be likely that this loss of interaction was dependent upon the phosphorylation state of Atg18. Such phosphorylation-dependent dissociation of Atg18 from the vacuole membrane was reported during hypoosmotic stress (Tamura *et al*., 2013). All these events together are likely to enhance the stringency of the system so that a subtle decrease in the levels of PtdIns(3,5)P_2_ causes rapid termination of PtdIns(3,5)P_2_-mediated downstream events. However, as previously stated by Tamura et al, the extremely low levels of PtdIns(3,5)P_2_, at ∼0.05% to 0.1% of total phosphoinositides under normal conditions, make it difficult to determine if lowering the levels of this lipid ligand or the modifications of Atg18 influencing its activity serve as the triggering event driving the release of Atg18 from the vacuole membrane. In addition to this, we observe that the localization of both Sch9 and Atg18 was PtdIns(3,5)P_2_-dependent, the 1-183 fragment of Sch9 can sense very minute depressions in the levels of PtdIns(3,5)P_2_ as compared to Atg18 under similar conditions (compare Figure 5E and 6C). This is because Sch9^1-183^ can exclusively bind to PtdIns(3,5)P_2_ whereas Atg18 can bind both PtdIns3P and PtdIns(3,5)P_2_ (Dove *et al*., 2004), the distribution of the former does not change upon ethanol treatment.

Our studies have shown that overexpressing Sec18 causes the restoration of growth defects of the temperature-sensitive mutants that cannot undergo homotypic vacuole fusion in the presence of ethanol (Figure 2). Recent reports show that Sec18 can bind to the tethering complex HOPS and enhance membrane fusion in the absence of Sec17 (Orr and Wickner, 2024). Consistently, increased ethanol tolerance through overexpression of Sec18 ties in with previous reports documenting the role of SM proteins (Vps33) and various subunits of the HOPS and CORVET complex all of which together are known to increase tolerance in the presence of ethanol (Teixeira *et al*., 2009). An avenue for future investigation into ethanol stress response would be to determine how the phosphorylation of Atg18 is regulated during this process and how the HOPS complex is modulated for the promotion of vacuole fusion. Changing how rapidly cells can accomplish homotypic vacuole fusion during ethanol stress can pave the way for augmenting ethanol yield using bioengineered yeast with better ethanol tolerance.

## MATERIALS AND METHODS

### Yeast strains

All yeast strains used for this study are listed in Table S1. Deletion strains are created by PCR-based homologous recombination of His3MX cassette (Longtine *et al*., 1998). The primers used for deletion PCR and deletion confirmation are listed in Table S2. The temperature-sensitive strains RSY269 (*sec17-1*) and RSY271 (*sec18-*1) were developed from the parent strain RSY255 (referred to as wild-type) (Novick *et al*., 1980; Kaiser and Schekman, 1990).

### Growth conditions

All the strains were grown at 28°C or as indicated otherwise. Cells were grown in YPAD medium containing 1%Yeast extract, 2% peptone, and 2% glucose. For live-cell imaging and spot assays cells were grown in Synthetic Dextrose Complete (SDC) medium or the same medium with the respective amino acid dropout (Burke *et al*., 2000).

### Stress conditions

For ethanol stress, liquid cultures were grown to mid-log phase to an optical density at OD_660_=0.6-0.8. Cells were harvested at 5000 rpm for 5 min and resuspended in SDC medium containing appropriate concentrations of ethanol (Merck) for the indicated time points. Cells were then washed twice with SDC and resuspended with the same medium containing ethanol. For spot dilution assay, ethanol was added into plates at desired concentrations. For salt and hypo-osmotic stress, liquid cultures grown to mid-log phase were harvested at 5000 rpm for 5 min and resuspended in Synthetic Complete (SC) medium supplemented with either 0.5 M NaCl or water respectively and incubated for indicated time points. Cells were then washed twice and resuspended in the same medium. For thermal stress, log phase cells were shifted to 37°C or 42°C for 1 h before visualization (Keuenhof *et al*., 2022).

### Plasmid construction

The constructs used in this study were generated by PCR amplifying the desired ORFs using gene-specific primers listed in Table S2 and then cloned under the control of either constitutive or inducible promoters. The clones were confirmed by DNA sequencing. The construct list is tabulated in Table S3.

### Live cell imaging

Overnight-grown cultures were diluted to an OD_660_ = 0.2 and were grown until the mid-log phase (OD_660_ = 0.6-0.8). Cells were harvested, and vacuoles were stained with either 7-amino-4chrolomethylcoumarin (CMAC) (Thermo Fisher Scientific) following manufacturer’s protocol or FM4-64 (Thermo Fisher Scientific) (Vida and Emr, 1995). For FM4-64 staining, mid-log phase cells were harvested and dissolved in YPD followed by the addition of 30 µm of FM4-64 at 1:50 dilution for 30 min. Cells were washed twice in SDC media and chased at 28°C for 40 min. Cells were washed twice in SDC medium followed by adding ethanol at the desired concentration in SDC media. The cells were washed following ethanol stress and resuspended in the same media containing ethanol. For luminal vacuole staining, mid-log phase cells were supplemented with 0.1 mM CMAC for 30 min followed by adding ethanol. To determine the vacuolar morphology of *sec17-1* and *sec18-1*, cells were initially grown at 24°C to mid-log phase. To induce temperature-sensitivity, cells were shifted to 26°C or 28°C for 1 h. Ethanol was added to attain a concentration of 10% for 30 min before visualization.

For probing the localization of PtdIns3P, two copies of the FYVE domain from the human Hrs protein were N-terminally tagged with GFP (*GFP-FYVE_2_*) (Zieger and Mayer, 2012). Sch9^1-183^ was C-terminally tagged with RFP and was used as a sensor for PtdIns(3,5)P_2_. To determine the localization of Atg18 post ethanol stress, yeast cells containing *pGAL-GFP-ATG18* in pRS315 were grown in Synthetic Dextrose Media without leucine in the presence of 2% raffinose at 28°C. Atg18 was expressed by inducing the GAL1 promoter with 2% galactose for 4-6h (Roy *et al*., 2015). Imaging was performed using a confocal laser scanning microscope (Stellaris, Leica Microsystems) with a 63X oil immersion objective having a numerical aperture of 1.4. Image acquisition was done using the Leica Application Suite X software and brightness and contrast were adjusted using Adobe Photoshop CS3 and assembled using Adobe Illustrator CS3.

### Image analysis

For image quantification, the percentage of cells carrying variations of vacuole lobes was counted using Image J (version 2.3.0) from three independent biological replicates, with approximately 60-80 cells per replicate (Grziwa *et al*., 2023). The mean and standard deviations were calculated and plotted using GraphPad Prism 10. For depth coding and three-dimensional image rendering of vacuolar invaginations, individual image *z-*stacks were recorded at 100-500 nm spanning the entire depth of the cells, and movies were created using the Leica Application Suite X software.

### Spot assay

Overnight cultures in appropriate SD medium were diluted to an O.D = 0.2 and grown to mid-log phase for OD_660_ = 0.6-0.8. Cells were diluted again to an O.D = 0.1 and were serially diluted (1:10) and spotted onto SD plates containing appropriate concentrations of ethanol.

### Yeast two-hybrid assay

The constructs were expressed in pGAD424 (*LEU2* selection marker) as fusions of either the Gal4-activation domain (AD) or in pGBT9 (*TRP1* selection marker) (Clonetech Laboratories) as fusions of the Gal4-DNA binding domain (BD). Binary constructs expressing AD or BD fusion proteins were cotransformed into PJ69-4A and incubated at 24°C. The transformants were selected on leucine tryptophan dropout plates. An equal number of cells were spotted on selective media to monitor interactions through this assay. The growth of the transformants was monitored on Synthetic minimal media lacking leucine tryptophan (LT^-^) either with or without ethanol. Positive interactions were screened by the activation of two downstream reporter genes, *HIS3* and *ADE2*, which was monitored by the growth on leucine tryptophan histidine dropout plates containing 2.5 mM 3-amino-triazole (LTH^-^3AT) and leucine tryptophan adenine (LTA^-^) dropout plates both in the presence and absence of ethanol Both LTH^-^3AT and LTA^-^ dropout plates were incubated at 24°C for 10 days (Jin *et al*., 2008; Lang *et al*., 2017).

## Acknowledgments

The authors acknowledge Prof. Alok Kumar Sil for his valuable suggestions during the study and for proofreading the manuscript. We thank, Prof. Randy Schekman, and Prof. Peter Novick for providing yeast strains. We also thank Prof. Christian Ungermann, Prof. Andreas Mayer, and Prof. Atin Kumar Mandal for sharing plasmids. We thank Dr. Shankari Prasad Datta and Dr. Ananya Jana for generating resources that helped in our study. The authors thank the Central Instrument Facility of Bose Institute for their contribution to this work, particularly The Confocal Imaging Facility and the DNA Sequencing Facility. We are grateful to Prantik Saha, Sheolee Ghosh Chakraborty, and Ayantrisha Biswas for imaging and Leica Microsystems for their technical assistance. We are thankful to the members of the Sarkar Laboratory for their discussions and for providing valuable comments.

## Competing interest

The authors declare no financial or competing interests.

## Author Contributions

Conceptualization: PM, SS; Methodology: PM, BH, SS; Software: PM; Validation: PM, SS; Formal Analysis: PM, BH, SS; Investigation: PM, BH; Resources: SS; Data curation: PM, BH, TG, NP, SS; Writing-Original draft: PM; Writing-review and editing: PM, SS; Visualization: PM, SS; Supervision: SS; Project administration: SS; Funding acquisition: SS

## Funding

This work was supported by intramural funds from Bose Institute, Kolkata (R/16/19/1620). Fellowship support: PM from CSIR (09/015(0545)/2019-EMR-I); BH from UGC (221610061747); TG from UGC (721/(CSIR-UGC NET DEC 2018)); NP from (09/015(0534)/2019-EMR-I)

## Abbreviations

PIKFYVE: phosphatidylinositol-3-phosphate-5 kinase
PtdIns: phosphatidylinositol
PtdIns3P: phosphatidylinositol-3-phosphate
PtdIns(3,5)P_2_: phosphatidylinositol-3,5-bisphosphate
SNARE: <u>s</u>oluble- *<u>N</u>*-ethylmaleimide-sensitive factor <u>a</u>ttachment protein <u>re</u>ceptor
NSF: *<u>N</u>*-ethylmaleimide-<u>s</u>ensitive <u>f</u>usion protein
SNAP: α-Soluble NSF attachment protein
FM4-64: (N-(3-Triethylammoniumpropyl)-4-(6-(4- (Diethylamino) Phenyl) Hexatrienyl) Pyridinium Dibromide)
3-AT: 3-amino triazole
AD: Gal4 Activation Domain
BD: Gal4 DNA Binding Domain

## Notes

### Competing Interest Statement

The authors have declared no competing interest.

