## Supplementary File for "Phosphatidylinositol-3,5-bisphosphate mediated vacuolar morphology modulation is integral to ethanol stress response"

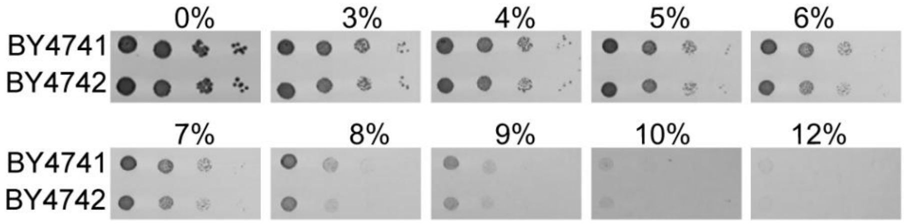


**Supplementary Figure S1.** Ethanol sensitivity of BY4741 and BY4742**.** (A) Cells were grown to mid-log phase and sequentially diluted 10-fold and spotted in Synthetic Complete Medium containing increasing concentrations of ethanol.

**Video 1 and 2. Ethanol-induced vacuolar invaginations.** Three-dimensional image rendering of individual *z*-stacks and depth coding to show the vacuolar membrane imagination from the surface of the vacuole (indicated in blue) to the lumen (indicated in red). Distance is in μm. The approximate depth of the invagination is ~2 μm.

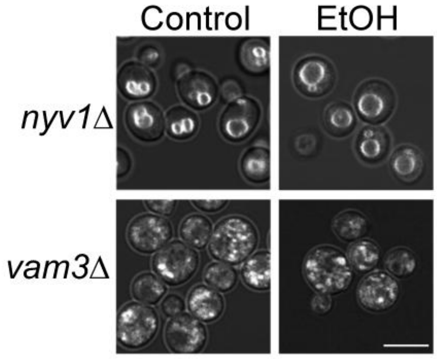


**Supplementary Figure S2**. Vacuole morphology of *nyv1*Δ and *vam3*Δ. Vacuolar membranes of *nyv1*Δ and *vam3*Δ were stained with FM4-64 and vacuolar morphology was examined following exposure to 10% ethanol for 30 min. Scale bar: 5 μm

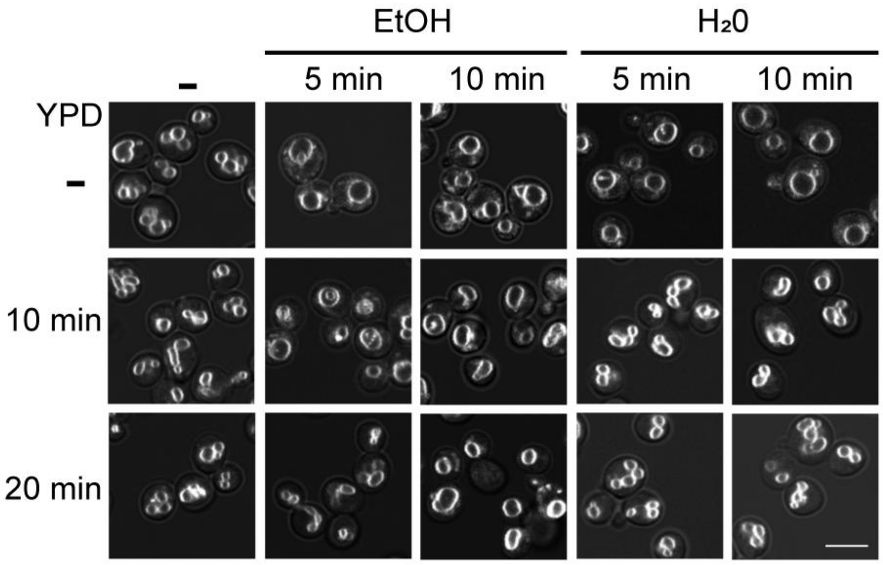


**Supplementary Figure S3**. Recovery of vacuole morphology following vacuole enlargement in the presence of ethanol**.** Vacuolar membranes of BY4742 were stained with FM4-64 and vacuolar morphology was examined for ethanol and hypoosmotic stress-treated cells for 5 min and 10 min followed by the addition of YPAD for either 10 min or 20 min. Scale bar: 5 µm.

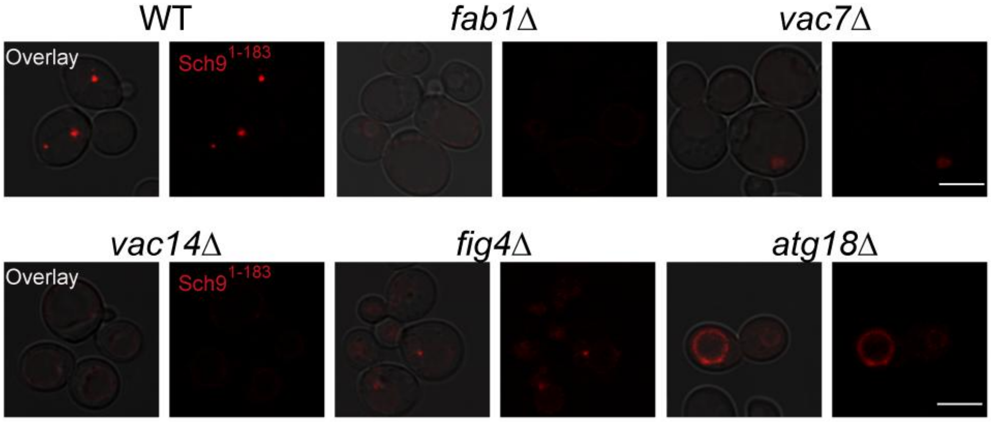


**Supplementary Figure S4**. **Sch9$^{1\text{-}183}$localization in mutants of the Fab1 complex.** Sch9$^{1\text{-}183}$ fragment was C-terminally tagged with RFP and was expressed under a constitutive promoter in wild-type, *fab1*Δ*, vac7*Δ*, vac14*Δ*, fig4*Δ*,* and *atg18*Δ cells. Scale bar: 5 µm

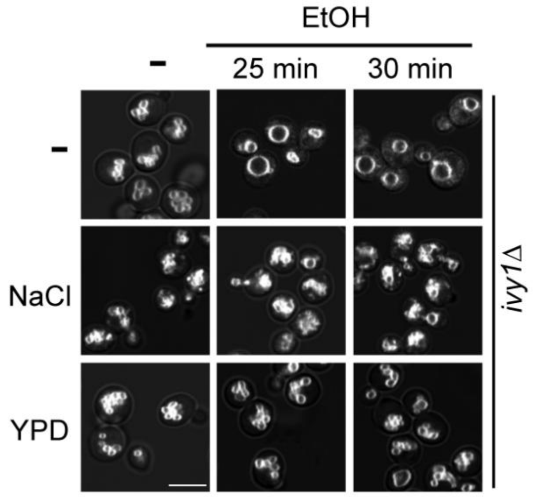


**Supplementary Figure S5.** Analysis of vacuole fission in ethanol-treated *ivy1*Δ cells. *ivy1*Δ were stained with FM4-64 and vacuolar morphology was examined for ethanol-treated cells for 25 min and 30 min followed by the addition of Nacl or YPAD for 10 min. Scale bar: 5 µm

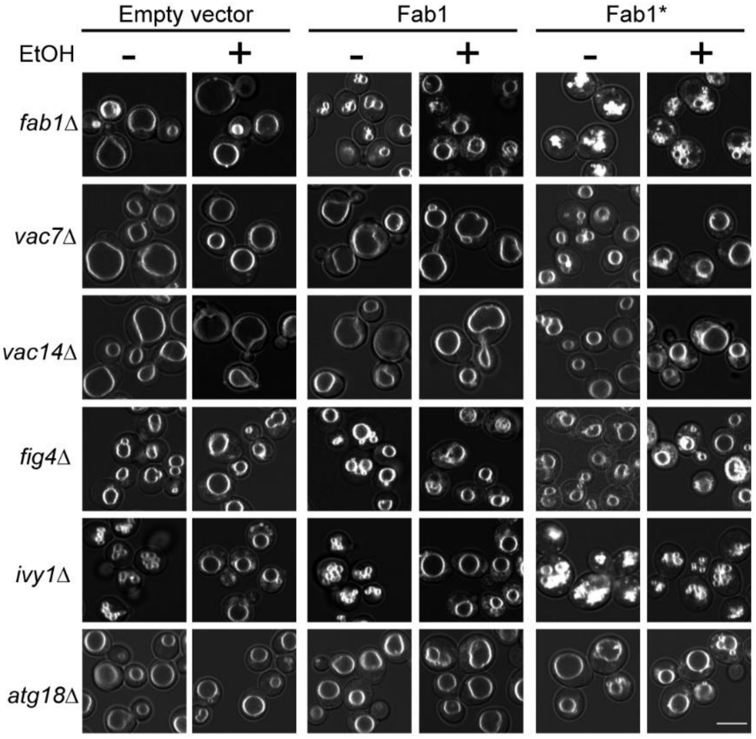


**Supplementary Figure S6.** Vacuolar morphology was examined for Fab1 complex mutants in the presence of ethanol**.** Vacuoles of mutants were stained with FM4-64 and was examined in the presence and absence of ethanol expressing either the empty vector (negative control), or wild-type copy of Fab1 or the Fab1 hyperactive allele (Fab1*). Scale bar: 5 µm.

**Table S1. List of strains used in this study**

| Sl. No. | Strain | Genotype | Reference |
|---|---|---|---|
| 1. | BY4742 | MATα *his3Δ1 leu2Δ0 lys2Δ0 ura3Δ0* | Euroscarf |
| 2. | PJ694-A | *MATa trp1-Δ901 leu2-3,112 901 ura3-52 his3-Δ200 gal4Δ gal8Δ GAL2-ADE2 LYS2::GAL1-HIS3 met2::GAL7-lacZ* | Clonetech laboratories |
| 3. | RSY249 | MATα *ura3-52 his4-619* | (Novick *et al.*, 1980) |
| 4. | RSY271 | MATα *ura3-52 his4-619 sec18-1* | (Novick *et al.*, 1980) |
| 5. | RSY269 | MATα *ura3-52 his4-619 sec17-1* | (Novick *et al.*, 1980) |
| 6. | *nyv1*Δ | BY4742, *NYV1::HIS5* | This study |
| 7. | *vam3*Δ | BY4742, *VAM3::HIS5* | This study |
| 8. | *ypt7*Δ | BY4742, *YPT7::HIS5* | This study |
| 9. | *vps34*Δ | BY4742, *VPS34::HIS5* | This study |
| 10. | *fab1*Δ | BY4742, *FAB1::HIS5* | This study |
| 11. | *vac7*Δ | BY4742, *VAC7::HIS5* | This study |
| 12. | *vac14*Δ | BY4742, *VAC14::HIS5* | This study |
| 13. | *fig4*Δ | BY4742, *FIG4::HIS5* | This study |
| 14. | *ivy1*Δ | BY4742, *IVY1::HIS5* | This study |
| 15. | *atg18*Δ | BY4742, *ATG18::HIS5* | This study |

| Table S2. Primer sequences used in this study | | |
|---|---|---|
| **Sl. No.** | **Primer sequence (5'-3')** | **Purpose** |
| 1. | ATTAAGCTGTTAGAGCATTGGACTTTTATATTTTTACC AACGGATCCCCGGGTTAATTAA | Forward primer for deletion of *Nvy1* |
| 2. | TGTTATTGTCGTGGGACAGCTCCCCTTTTTTTTTATT TACGAATTCGAGCTCGTTTAAAC | Reverse primer for deletion of *Nvy1* |
| 3. | GCTGTTAGAGCATTGGACTT | Forward primer for *Nyv1* deletion confirmation |
| 4. | GTACACAGTTTTGGGGTCA | Reverse primer for *Nyv1* deletion confirmation |
| 5. | CAAATTGGCCAACTAATATCCACTGCAGAAAGTTGA GATTCGGATCCCCGGGTTAATTAA | Forward primer for deletion of *Vam3* |
| 6. | TCTACCAGAAAGTCTGTGCAATGCGCGTTTAAGGAG ATTAGAATTCGAGCTCGTTTAAAC | Reverse primer for deletion of Vam3 |
| 7. | GCCAACTAATATCCACTGCA | Forward primer for Vam3 deletion confirmation |
| 8. | GCATTGACTTCTTGTACCGC | Reverse primer for *Vam3* deletion confirmation |
| 9. | TCCATATAGAAACCCCTTCTGTATCAATTCAAATTAA GTGCGGATCCCCGGGTTAATTAA | Forward primer for deletion of *Ypt7* |
| 10. | AAAGGATTACATAATAGAAGATACAATTAAGTAGTA CAGCGAATTCGAGCTCGTTTAAAC | Forward primer for deletion of *Ypt7* |
| 11. | CCACTTCTTATCCATATAGAAACCC | Forward primer for *Ypt7* deletion confirmation |
| 12. | GAAATTCATCTCGCCAAGAC | Reverse primer for *Ypt7* deletion confirmation |
| 13. | ACT**GGATCC**GTAATGATAAGAAGTGGGCAGCACCG | Forward primer for cloning *Sec17* for complementation |
| 14. | ATCTA**GTCGAC**GATGGCTTCAACTTCGCTCTTGC | Reverse primer for cloning *Sec17* for complementation |
| 15. | AAT**GGATCC**TCGGTATCACACTATACGTTAGGCG | Forward primer for cloning *Sec18* for complementation |
| 16. | TACGT**GTCGAC**TGACAGGCAGATTCCAATGACAG | Reverse primer for cloning *Sec18* for complementation |
| 17. | AAG**GTCGAC**ATGAGCGAGCTGATC | Forward primer for cloning RFP in pRS45 |
| 18. | CCA**CTGCAG**TTATCTGTGCCCCA | Reverse primer for cloning RFP in pRS45 |
| 19. | GGC**GAATTC** ATACGAATATACAATATGATG | Forward primer for cloning $Sch9^{1-183}$ for PI(3,5)$P_2$ localization |
| 20. | ATA**GGATCC**TTTACCCCTAGGAATATCGGTA | Reverse primer for cloning $Sch9^{1-183}$ for PI(3,5)$P_2$ localization |

| | | |
|---|---|---|
| 21. | CATCTCCGTGAAGCATTGAGGGAAGGGTTTAACTCC AACACGGATCCCCGGGTTAATTAA | Forward primer for deletion of *Vps34* |
| 22. | GTGACGAAATTTAAATTTTGAAGCACCAATTATCAA CCAA GAATTCGAGCTCGTTTAAAC | Reverse primer for deletion of *Vps34* |
| 23. | GAAGCATTGAGGGAAGGGTTT | Forward primer for *Vps34* deletion confirmation |
| 24. | GCTTAGTCCAGGATTGTTCTTC | Reverse primer for *Vps34* deletion confirmation |
| 25. | AGGTAGCTTCCATCCTGTACATGCAAGACCGTCACA CAGCCGGATCCCCGGGTTAATTAA | Forward primer for deletion of *Fab1* |
| 26. | AAAAAAAAGTTACAGAATATAACTTGTACACGTTTA TGTAGAATTCGAGCTCGTTTAAAC | Reverse primer for deletion of *Fab1* |
| 27. | GCTTCCATCCTGTACATGCAA | Forward primer for *Fab1* deletion confirmation |
| 28. | TTTAATAGTTCTAACGCGTACCCAC | Reverse primer for *Fab1* deletion confirmation |
| 29. | TTCATCTCAGGCAAGTTAAAGCATTTGGGAAACGTG CTAGCGGATCCCCGGGTTAATTAA | Forward primer for deletion of *Vac7* |
| 30. | AAAATACCCAGCTTTGACGAAAAAGCTACATTCTTA ACACGAATTCGAGCTCGTTTAAAC | Reverse primer for deletion of *Vac7* |
| 31. | GCATTTGGGAAACGTGCTA | Forward primer for *Vac7* deletion confirmation |
| 32. | CAAGTCGGCGTTAGAATTC | Reverse primer for *Vac7* deletion confirmation |
| 33. | TGCTGTGCTTATCTGCTCAGGCTACAACAGGAACTG GAACCGGATCCCCGGGTTAATTAA | Forward primer for deletion of *Vac14* |
| 34. | ATTTCTTAACCAAAGATGCTTTCAATCAGGTAATGG GTAGGAATTCGAGCTCGTTTAAAC | Reverse primer for deletion of *Vac14* |
| 35. | CTTTTGATGCTGCTGTGCTTA | Forward primer for *Vac14* deletion confirmation |
| 36. | TCTTACTTGGTCATTCTGGTCG | Reverse primer for *Vac14* deletion confirmation |
| 37. | AAAATATTGTTAAAGAAAGTAACAGGAAGAGAAAT CGGATCGGATCCCCGGGTTAATTAA | Forward primer for deletion of *Ivy1* |
| 38. | CTCCATTTCTATATAAAAAGCATACATAGAGTTACAA ATTGAATTCGAGCTCGTTTAAAC | Reverse primer for deletion of *Ivy1* |
| 39. | GTAACAGGAAGAGAAATCGG | Forward primer for *Ivy1* deletion confirmation |
| 40. | ATCATTGCAACCCTTCAAC | Reverse primer for *Ivy1* deletion confirmation |
| 41. | GTTTTGAAACGTAGTAACGTAACGCAAAGCAAAAA AGAAACGGATCCCCGGGTTAATTAA | Forward primer for deletion of *Fig4* |
| 42. | TGACCGAATATTTAAATTTCTATTTAAGAGTCATATAA ATGAATTCGAGCTCGTTTAAAC | Reverse primer for deletion of *Fig4* |
| 43. | GTAGTAACGTAACGCAAAGCA | Forward primer for *Fig4* deletion confirmation |
| 44. | AATGAATCCCAGAAGACCGT | Reverse primer for *Fig4* deletion confirmation |
| 45. | TTCCAGTTAACTCTGTATCCTTTTCTTCTTCGGCCTG ACACGGATCCCCGGGTTAATTAA | Forward primer for deletion of *Atg18* |

| | | |
|---|---|---|
| 46. | CGTTGTGACGTACGGAAGGCAGCGCGAGACACTTC CGTGAGAATTCGAGCTCGTTTAAAC | Reverse primer for deletion of *Atg18* |
| 47. | CTTTTCTTCTTCGGCCTGA | Forward primer for *Atg18* deletion confirmation |
| 48. | GATCCAAGTCGCTAATGTCAC | Reverse primer for *Atg18* deletion confirmation |
| 49. | AGA**GAATTC**ATGGAAAAATCGATTGCCAAAG | Forward primer for cloning *Vac14* for Y2H |
| 50. | CGG**CTGCAG**TTATTTTTTAATTTATCGGATAC | Reverse primer for cloning *Vac14* for Y2H |
| 51. | GGC**GGATCC**AGATGACAGAAGAAGATAGAAAGC | Forward primer for cloning *Vac7* for Y2H |
| 52. | AAT**GTCGAC**TCACTTCTTACCAGGATGGACC | Reverse primer for cloning *Vac7* for Y2H |
| 53. | AGA**GGATCC**AGATGAACAATGATGCAATGG | Forward primer for cloning *Fig4* for Y2H |
| 54. | GCG**GTCGAC**TTATTGAAAATCAAGTTGTATATC | Reverse primer for cloning *Fig4* for Y2H |
| 55. | AAC**GGATCC**AGATGCCTGACAATAATACGG | Forward primer for cloning *Ivy1* for Y2H |
| 56. | GGC**GTCGAC**TTATATATTACTTGACTGGTCTTGC | Reverse primer for cloning *Ivy1* for Y2H |
| 57. | GGC**GAATTC**ATGTCTGATTCATCACCTACTA | Forward primer for cloning *Atg18* for Y2H |
| 58. | GAG**GTCGAC** TCAATCCATCAAGATGGAATAC | Reverse primer for cloning *Atg18* for Y2H |
| 59. | AAT**CCGCGG**TGTCTGATTCATCACCTAC | Forward primer for cloning *Atg18* |
| 60. | GAC**GAGCTC**TCAATCCATCAAGATGGAATAC | Reverse primer for cloning *Atg18* |

Note: The restriction sites used in the primers are underlined

| Table S3. List of constructs used in this study | | | |
|---|---|---|---|
| **Sl. No.** | **Cloned gene** | **Description** | **Primers used** |
| 1. | *SEC17* | p(PYK1)-*SEC17* (pRS425 *SEC17 LEU2*) | 13 and 14 |
| 2. | *SEC18* | p(PYK1)-*SEC17* (pRS426 *SEC18 URA3*) | 15 and 16 |
| 3. | *RFP* | p(PYK1)-*RFP* (pRS425 *RFP LEU2*) | 17 and 18 |
| 4. | $SCH9^{1\text{-}183}$ | p(PYK1)- $SCH9^{1\text{-}183}$-*RFP* (pRS425 $SCH9^{1\text{-}183}$-*RFP LEU2*) | 19 and 20 |
| 5. | *FYVE* | $FYVE_2$-*GFP* | (Zieger and Mayer, 2012) |
| 6. | *Fab1** | *Fab1-5* (Hyperactive mutant of Fab1) | (Malia *et al.*, 2018) |
| 7. | *Fab1* | *Fab1-WT* | A kind gift from Prof. Christian Ungermann |
| 8. | *VAC14* | p(ADH1)-GAL4AD-*VAC14* (pGAD424 *VAC14 LEU2*) | 49 and 50 |
| 9. | | p(ADH1)-GAL4BD-*VAC14* (pGBT9 *VAC14 TRP1*) | |
| 10. | *VAC7* | p(ADH1)-GAL4AD-*VAC7* (pGAD424 *VAC7 LEU2*) | 51 and 52 |
| 11. | | p(ADH1)-GAL4BD-*VAC7* (pGBT9 *VAC7 TRP1*) | |
| 12. | *FIG4* | p(ADH1)-GAL4AD-*VAC7* (pGAD424 *FIG4 LEU2*) | 53 and 54 |
| 13. | | p(ADH1)-GAL4BD-*VAC7* (pGBT9 *FIG4 TRP1*) | |
| 14. | *IVY1* | p(ADH1)-GAL4AD-*VAC7* (pGAD424 *IVY1 LEU2*) | 55 and 56 |
| 15. | | p(ADH1)-GAL4BD-*VAC7* (pGBT9 *IVY1 TRP1*) | |
| 16. | *ATG18* | p(ADH1)-GAL4AD-*VAC7* (pGAD424 *ATG18 LEU2*) | 57 and 58 |
| 17. | | p(ADH1)-GAL4BD-*VAC7* (pGBT9 *ATG18 TRP1*) | |
| 18. | *ATG18* | p(GAL1)-*GFP-ATG18* (pRS315 *GAL-GFP-ATG18*) | 59 and 60 |
